# Cryo-EM structure of the sub-50 kD PDZ-RhoGEF catalytic core–RhoA complex reveals an allosteric mechanism for GEF switch

**DOI:** 10.64898/2026.10.21.760610

**Authors:** Matthew Villanueva Braga, Emma Barpal, Sheng Li, Jennifer N. Cash

**Affiliations:** Department of Molecular and Cellular Biology, University of California – Davis; Davis, CA, USA; Department of Medicine, University of California – San Diego; La Jolla, CA, USA

## Abstract

PDZ-RhoGEF (PRG) drives cellular processes including migration through activation of the GTPase RhoA. The GEF switch (GS) is an accessory region N-terminal to the PRG catalytic core that significantly enhances its activity against RhoA, but its mechanism is unknown. We take an integrative structural biology approach, including cryogenic electron microscopy, to deduce the GS mechanism. Enzymatic assays define interactions between the GS and PRG catalytic core that are required for enhancement of activity. While SEC-SAXS data suggest that the GS does not alter the global structure of the catalytic core–RhoA complex, HDX-MS data reveal that the GS induces widespread stabilization, particularly in the RhoA switch regions. We determined a sub-4 Å resolution structure of 46 kD of the complex between the catalytic core and RhoA, revealing that Switch I is more flexible without the GS. Collectively, these data suggest both direct and allosteric factors in the mechanism of the GS.

## Introduction

The Dbl superfamily Rho guanine-nucleotide exchange factors (Dbl RhoGEFs) activate small Rho GTPases by catalyzing guanine nucleotide exchange to promote downstream cellular signaling and mediate critical processes including cell proliferation and migration^1,2^. Dbl RhoGEFs activate Rho GTPases through their conserved catalytic cores, composed of a Dbl homology (DH) domain almost invariably followed by a pleckstrin homology (PH) domain, which are subject to complex regulation by accessory regions outside of the core that vary widely across the 70 family members^2^. Unlike the well-studied catalytic core, our understanding of the precise mechanisms by which these accessory regions exert regulation on each RhoGEF are less understood. One such RhoGEF is PDZ-RhoGEF (PRG), which shows ubiquitous expression that is particularly strong in the brain, heart, placenta, ovaries, and testes and weaker in the prostate, colon, and lungs^3^. Outside of its normal functioning in cell migration^4,5^, PRG drives invasiveness and metastasis in various cancers^6–10^ and exacerbates other pathologies, including diabetes, obesity, cardiac hypertrophy, and bipolar disorder^11–13^, making it an attractive therapeutic target. Mice deficient in PRG do not show obvious adverse phenotypes, further supporting PRG inhibition as a viable therapeutic strategy^14^. However, the rational design of therapeutics targeting PRG requires an understanding of the molecular mechanisms regulating it, which remain largely unclear. To date, there are few drugs that have been developed to inhibit RhoGEFs^3^, but understanding accessory region-mediated regulatory mechanisms, which are unique between individual RhoGEFs^2^, will enable us to pinpoint sites on the proteins that could serve as feasible targets for selective inhibitors.

PRG specifically activates the GTPase RhoA downstream of G protein-coupled receptor (GPCR) signaling, resulting in actin cytoskeletal rearrangement and cell migration^15,16^. Within the catalytic core of PRG, the DH domain directly engages and stabilizes RhoA Switch I (residues 27-43) and Switch II (residues 57-78) to accelerate the release of GDP, forming a reaction intermediate complex between PRG and nucleotide-free RhoA. GTP, which is more intracellularly abundant than GDP, then binds to RhoA and displaces PRG^2,17,18^. The PH domain associates with the DH domain and RhoA to form a minimal buried surface, enhancing the activity of the DH domain by 30-fold^16^. PRG accessory regions include the PSD95, dis cs large, and zonula occludens-1 (PDZ) domain, the regulators of G protein signaling homology (RH) domain, and the GEF switch (GS) (Figure 1A)^2,3,19–21^. The GS is a 17-residue accessory region N-terminal to the catalytic core of PRG^22,23^. Up to a third of Dbl RhoGEFs, most of which show a capacity to activate RhoA, could harbor this accessory region, based on sequence analysis (Figure S1). The GS enhances activation of RhoA in p115-RhoGEF and leukemia-associated RhoGEF (LARG), two GS-containing RhoGEFs that are closely related to PRG^22,23^. Whether the GS serves this same function in PRG is unknown. Multiple crystal structures of the DH/PH catalytic core plus GS exist for these three RhoGEFs in complex with RhoA (GS-DH/PH–RhoA), showing that the GS buries surface area on both the DH domain and RhoA Switch I^16,22,23^. Yet, structural data on a complex without the GS is lacking, hindering the determination of its GEF-enhancing mechanism.

**Figure 1.**
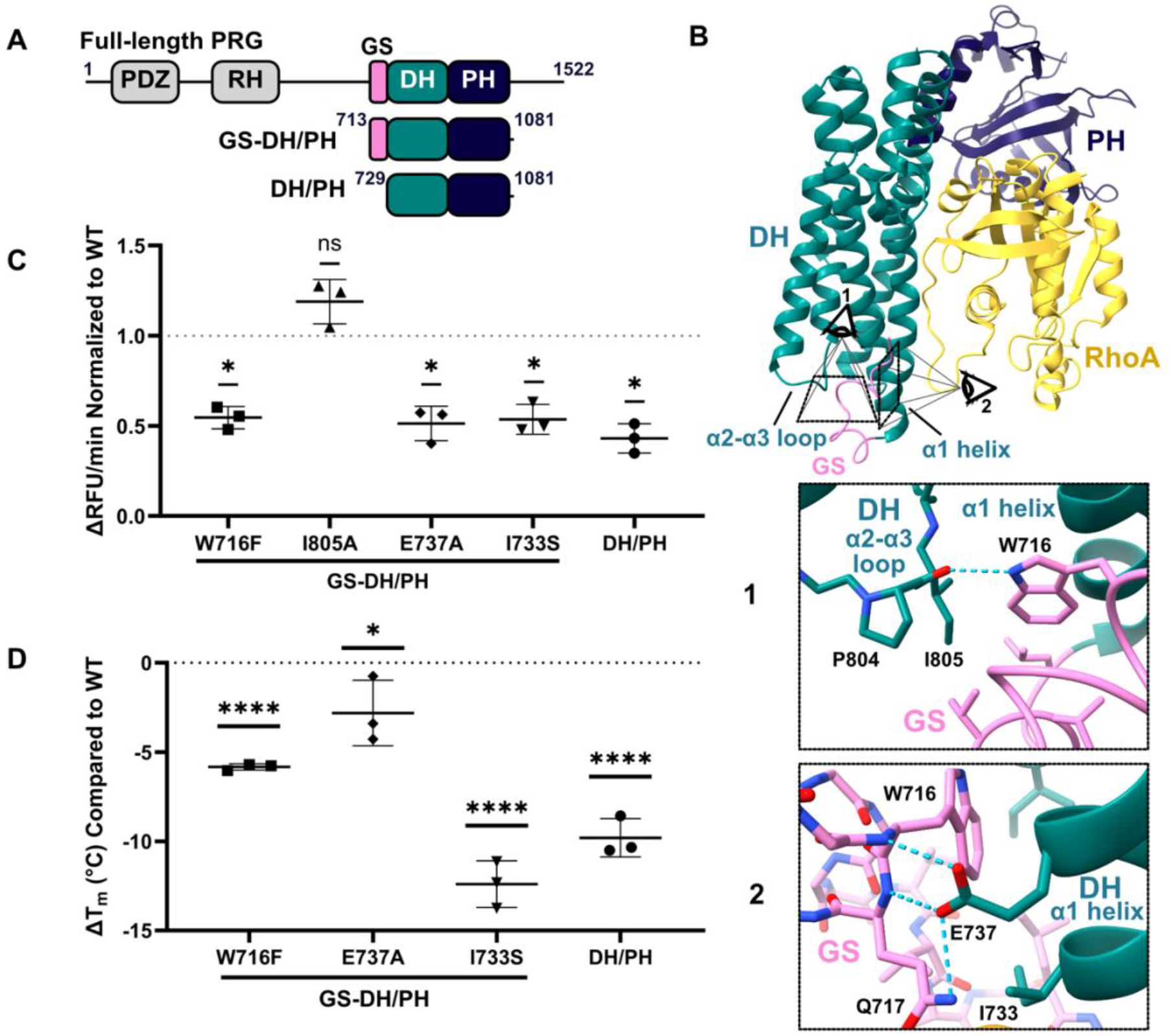
The GEF switch enhances PRG activity against RhoA through interaction with the DH domain. A) Domain layout of PRG constructs. B) GS-DH/PH–RhoA crystal structure 5JHG showing the GS-DH interface highlighting the interactions between the GS and DH α2-α3 loop (1) and DH α1 helix (2). Trp716 from the GS stabilizes interactions with the α1 helix and α2-α3 loop of the DH domain through both hydrogen bonding and the hydrophobic effect. E737 of the α1 helix contributes additional hydrogen bonds with Q717 of the GS. C) mant-GTP exchange assay data showing the activities of purified PRG GS-DH/PH variants against RhoA normalized to WT GS-DH/PH. Individual points represent biological replicates as an average of three technical replicates each (Figure S4A). Statistical significance was assessed using a one-sample t-test, comparing the relative activity of each mutant to the WT value of 1. A Holm-Šídák correction was applied to account for multiple comparisons (* = p < 0.05). D) DSF data comparing the melting temperatures of PRG GS-DH/PH variants as a change from WT. Individual points represent biological replicates as an average of three technical replicates each (Figure S4B). DH/PH represents data presented in Anderson et al^40^. Statistical significance was assessed using a one-way ANOVA followed by Dunnett’s correction for multiple comparisons (* = p < 0.05, **** = p < 0.001).

Here, we show that the GS enhances guanine nucleotide exchange in PRG, similar to LARG and p115-RhoGEF. We compare the complex comprising the catalytic core of PRG and nucleotide-free RhoA in the presence and absence of the GS to investigate the mechanism by which it enhances PRG activity against RhoA using an integrative structural biology approach including size-exclusion chromatography small-angle X-ray scattering (SEC-SAXS), hydrogen-deuterium exchange mass spectrometry (HDX-MS), and cryogenic electron microscopy (cryo-EM), complemented by enzymatic assays. These complexes are challenging cryo-EM targets, as they approach the theoretical lower size limit of proteins resolvable by cryo-EM and present unique challenges associated with the exceptionally low signal-to-noise ratio of particularly small particles^24,25^. We present the optimization of our cryo-EM workflow required for this small sample, leading to a sub-4 Å resolution reconstruction representing ∼46kD of the DH/PH– RhoA complex and to a low-resolution reconstruction of the ∼65kD GS-DH/PH–RhoA complex. These integrative structural biology data collectively reveal that the GS allosterically stabilizes the PRG DH and PH domains and RhoA switch regions within the PRG– RhoA guanine nucleotide exchange intermediate complex to enhance RhoA activation. To our knowledge, this is the first demonstration of a GS mechanism, notably an allosteric one, explaining its enhancement of GEF activity.

## Results

### A region N-terminal to the catalytic core enhances the GEF activity of PRG

Previous biochemical experiments showed that a region N-terminal to the canonical DH/PH catalytic tandem in both LARG^22^ and p115-RhoGEF^23^ potentiates activation of RhoA by each of the proteins. To investigate if this 17-amino acid region, coined the “GEF switch” (GS), exerts the same effects in PRG, we expressed and purified the DH/PH catalytic tandem with (residues 713-1081) and without (residues 729-1081) the GS (Figures 1A, S2A, and B). We observed that both PRG fragments eluted as single, narrow peaks at the expected volume in size-exclusion chromatography (SEC), indicating that the proteins are monodisperse and non-aggregated (Figure S2A and B). We used differential scanning fluorimetry (DSF) to obtain melting curves for both fragments from which we calculated melting temperatures (T_m_) of 44.3 °C and 34.9 °C for GS-DH/PH and DH/PH, respectively. This indicates that they are folded and that the GS provides a substantial stabilizing effect to the isolated DH/PH (Supplemental Table 1 and Figure S3). We then measured the activities of the purified PRG fragments against RhoA *in vitro* using a mant-GTP exchange assay that utilizes a fluorescent, nonhydrolyzable analog of GTP to produce a FRET-based signal that allows measurement of guanine nucleotide exchange in real-time. We observed an approximate twofold increase in the activity of PRG GS-DH/PH as compared to GS-DH/PH (Figures 1C). Taken together with the increased thermostability of GS-DH/PH (Figure 1D), these findings show that the GS enhances PRG-mediated guanine nucleotide exchange, potentially through stabilization of the catalytic core.

**Table 1.** SEC-SAXS Structural parameters.

|  | PRG DH/PH-RhoA | PRG GS-DH/PH-RhoA |
| --- | --- | --- |
| **Guinier analysis** |  |  |
| $I(0)$ (cm<sup>-1</sup>) | 45.29 +/- 0.12 | 45.24 +/- 0.2 |
| $R_g$ (Å) | 27.52 +/- 0.1 | 27.67 +/- 0.26 |
| $q_{min}$ (Å<sup>-1</sup>) | 0.0169 | 0.0149 |
| $qR_g$ max | 1.313 | 1.32 |
| Coefficient of correlation, $R^2$ | 0.997 | 0.996 |
| **Molecular weight analysis** |  |  |
| $M$ (Da) from $I(0)$ (ratio to expected value) | 6.09E4 (0.965) | 6.11E4 (0.942) |
| Porod volume (Å<sup>-3</sup>)<br>(ratio $V_p$/calculated $M$) | 8.13E4 (1.33) | 8.16E4 (1.34) |
| $M$ (Da) using Bayesian method<br>(ratio of $M$ to expected value) | 5.95E4 (0.943) | 5.81E4 (0.900) |
| **$P(r)$ analysis** |  |  |
| $I(0)$ (cm<sup>-1</sup>) | 45.32 +/- 0.09 | 45.17 +/- 0.09 |
| $R_g$ (Å) | 27.48 +/- 0.05 | 27.53 +/- 0.07 |
| $d_{max}$ (Å) | 85.0 | 88.0 |
| $q$ range (Å<sup>-1</sup>) | 0.0169 - 0.4702 | 0.0149 - 0.0477 |
| $\chi^2$ (total estimate from *GNOM*) | 0.788 | 0.724 |

Small-angle X-ray scattering (SEC-SAXS), pair distance distribution [ $P(r)$ ], radius of gyration ( $R_g$ ), Intensity at zero scattering angle [ $I(0)$ ], maximum particle diameter ( $d_{max}$ ), $M$ (molecular weight), scattering angle ( $q$ )

### The GS enhances PRG GEF activity in part through its interactions with the DH domain

To begin to investigate the mechanism behind this activity-enhancing effect of the GS, we analyzed the crystal structure of GS-DH/PH–RhoA. The tripartite hydrophobic pocket formed by the α1 helix and α2-α3 loop of the DH domain and the GS centers on Trp716, a component of the GS that is conserved among GS-containing RhoGEFs^22^. Trp716 packs with hydrophobic residues in the pocket and forms a conserved hydrogen bond with the main chain carbonyl of Pro804 in the α2-α3 loop (Figure 1B). To selectively assess the importance of this hydrogen bond to the GS mechanism in enhancing PRG activity, we mutated Trp716 in GS-DH/PH to phenylalanine. We reasoned that the bulky hydrophobic phenylalanine would stabilize the hydrophobic pocket while disrupting the hydrogen bond between the GS and the α2-α3 loop. GS-DH/PH W716F exhibited a twofold decrease in GEF activity against RhoA and a concomitant decrease in thermostability of approximately 6 °C as compared to GS-DH/PH WT, suggesting that GS function depends in part on this hydrogen bond with the α2-α3 loop (Figures 1C and D). GS-DH/PH W716F was still more thermostable than DH/PH, providing evidence that the structure of the GS is still at least partially intact in this variant (Figure 1D). In contrast, an I805A mutation targeting an α2-α3 loop hydrophobic residue that extends into the hydrophobic pocket did not significantly alter activity of GS-DH/PH. Thus, the hydrophobic packing contribution from the α2-α3 loop may not be as crucial for GS function as the hydrogen bond it forms with Trp716.

To test the contribution of the GS interaction with the α1 helix to its ability to enhance PRG activity, we generated two GS-DH/PH mutants targeting α1 helix residues Glu737 and Ile733 that pack against the GS. Residue Glu737 forms hydrogen bonds with the Trp716 amide and Gln717 sidechain and amide (Figure 1B). Variant E737A showed half of the activity of WT, indicating that Glu737 functions as a key determinant for GS function (Figures 1C). We chose to mutate Ile733 to serine to disrupt the hydrophobic packing of the α1 helix against the GS. Variant I733S displayed diminished thermostability by 12 °C, falling below the T_m_ of DH/PH, and half the activity of WT (Figures 1C and D). Thus, this hydrophobic contribution from the α1 helix is important for both the structure and function of the GS. To note, all variants that displayed decreased activity as compa red to WT produced S-shaped sigmoidal DSF melting curves characteristic of folded and non-aggregated proteins (Figure S3). Altogether, this supports that the GS enhances PRG activity against RhoA through multivalent interactions with the α2-α3 loop and α1 helix regions of the catalytic DH domain.

### PRG DH/PH– and GS-DH/PH–RhoA complexes are biochemically well-behaved and exhibit similar global structures

We next sought to determine if the GS alters the global structure of the PRG DH/PH–RhoA complex to carry out its activity enhancing function. We formed and purified complexes of PRG DH/PH and GS-DH/PH with nucleotide-free RhoA to high purity and yield (Figure S2C and D) and then obtained low resolution structural information on both complexes in solution using SEC coupled with small-angle X-ray scattering (SAXS) (Figure 2B and Table 1). Both DH/PH–RhoA and GS-DH/PH–RhoA eluted as single peaks with molecular weights corresponding to 1:1 PRG–RhoA complexes and Guinier plots showing high degrees of linearity, altoge ther indicating monodisperse, non-aggregated protein complexes (Figures 2A, C and D and Table 1). The dimensionless Kratky plot approximated a bell curve, suggesting that both complexes are globular as expected (Figure 2E). The scattering profiles of the two complexes shared a high degree of overlap and their radii of gyration were similar, reflecting structural similarity of the two complexes at low resolution (Figure 2B and Table 1). Altogether, the SEC-SAXS data indicate that the GS likely does not induce large-scale structural changes. The two complexes slightly differed in their pairwise distance distribution functions, histograms of distances between electron pairs within the protein, pointing to more subtle and regional GS-induced structural changes (Figure 2F).

**Figure 2.**
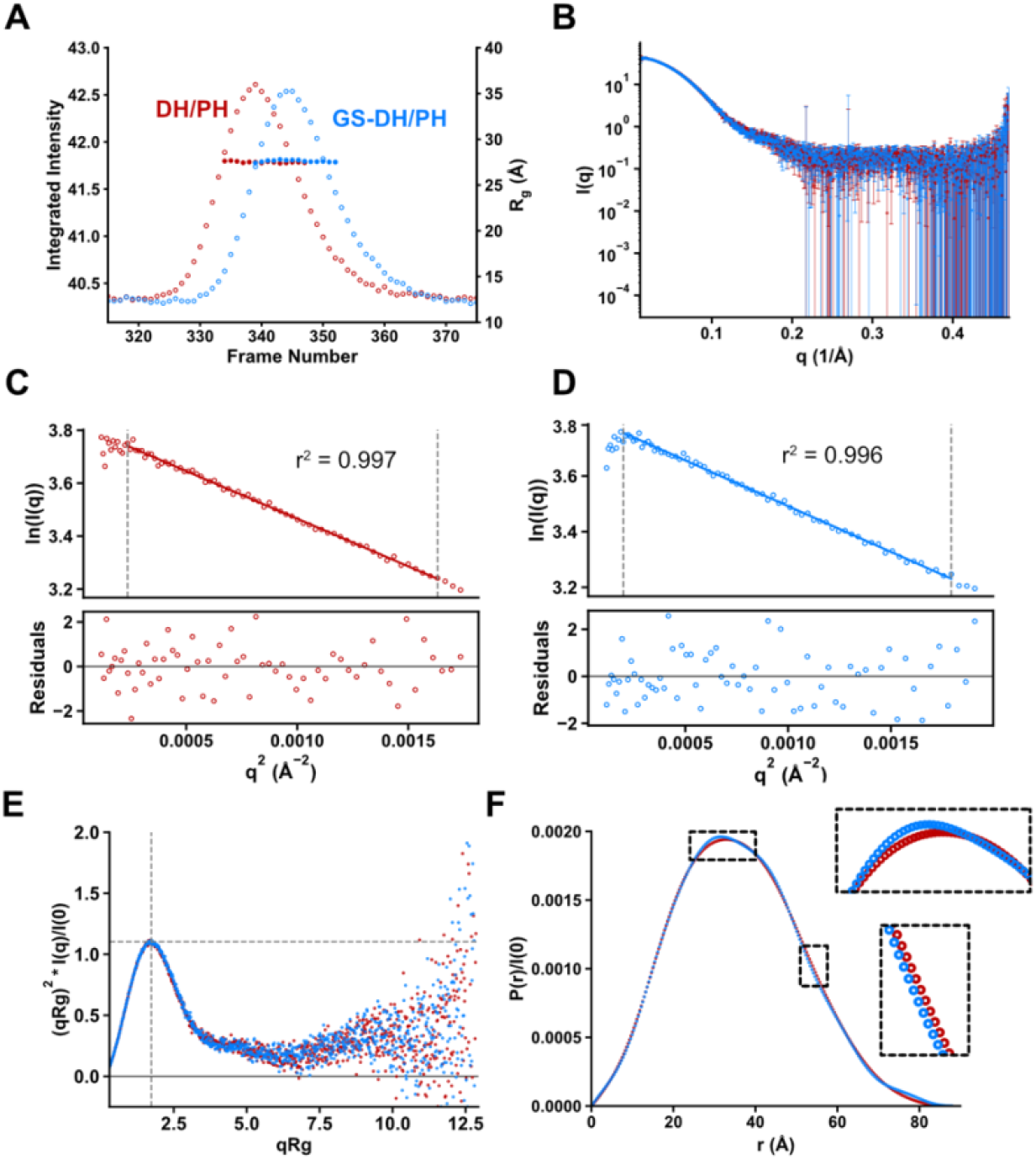
Structural characterization of PRG DH/PH–RhoA (red) and GS-DH/PH–RhoA (blue) samples using SEC-SAXS. A) SEC traces of DH/PH–RhoA and GS-DH/PH–RhoA with radius of gyration (Rg) plotted for fractions that correspond to the protein sample. Peaks and flat Rg’s across each peak indicate a monodisperse sample. B) Scattering intensities of DH/PH–RhoA and GS-DH/PH–RhoA at different spatial frequencies, q. Error bars represent error in intensity at each spatial frequency based on the counts in all of the pixels in a given angular band of the pixel array detector. High degree of overlap between the two profiles indicates global structural similarity. C and D) Guinier plots for DH/PH–RhoA and GS-DH/PH–RhoA. Residuals are plotted under the Guinier plots. High degree of linearity and flat residuals indicate monodispersity and lack of aggregation. E) Dimensionless Kratky plot for DH/PH–RhoA and GS-DH/PH–RhoA. A bell-shaped curve that peaks where the grey lines intersect indicates a globular system. F) Pair distance distribution function, or P(r), for DH/PH–RhoA and GS-DH/PH–RhoA. Boxed regions depict segments where the pair distance distribution functions differ between DH/PH–RhoA and GS-DH/PH– RhoA.

### HDX-MS shows that the GS increases protection of the DH domain and Switch regions of RhoA

To identify potential regional structural changes, we obtained hydrogen-deuterium exchange mass spectrometry (HDX-MS) profiles of DH/PH–RhoA and GS-DH/PH–RhoA complexes. In HDX-MS, proteins are incubated in a solution containing deuterated water and then quenched, proteolyzed, and analyzed through mass spectrometry to measure deuterium incorporation across different regions of the proteins. Comparison of these profiles between DH/PH–RhoA and GS-DH/PH–RhoA revealed widespread GS-induced decreases in deuterium incorporation in both PRG and RhoA (Figures 3A). Within the DH domain, the N-terminus of the α1 helix showed a substantial 47% decrease in deuteration, while the region around the α2-α3 loop showed a 12% decrease (on average after 10,000 s) in the presence of the GS as compared to the absence (Figures 3A and B). These results support that the GS interacts with these proximal regions to execute its GEF enhancing function, consistent with our activity and thermostability assay data (Figures 1C and D). Within RhoA, residues 25-84 exhibited greatly decreased deuteration in the presence of the GS (Figure 3A). These span the switch regions of RhoA, Switch I (residues 27-43) and Switch II (residues 57-78)^15^, critical for guanine nucleotide binding and effector recognition^16^ (Figure 3B). In the crystal structures of GS-DH/PH–RhoA^23^, Switch I lies proximal to the GS with a small buried surface of ∼100 Å^2^, while also engaging the N-terminus of DH α1 helix (Figure 3B). In the presence of the GS, Switch I deuteration decreased by 11%, suggesting stabilization by the GS. Surprisingly, Switch II deuteration decreased by 18%, more drastically than Switch I, despite being located 36 Å away from the GS, providing evidence for an allosteric stabilization mechanism (Figure 3B). Altogether, the direct and allosteric stabilization of these regions of PRG and RhoA by the GS could underlie the mechanism by which it enhances PRG activity.

**Figure 3.**
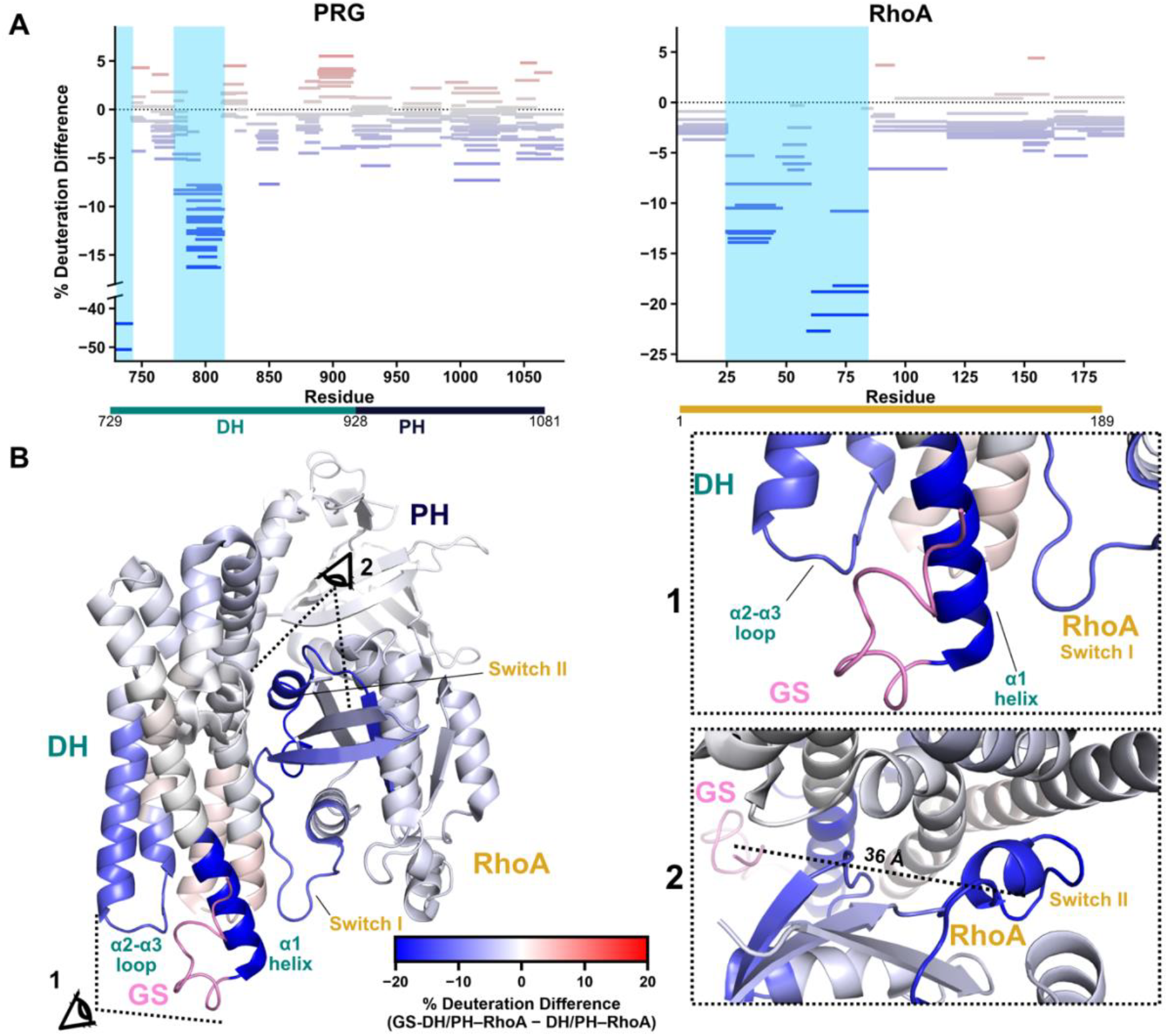
The GEF Switch allosterically stabilizes the switch regions of RhoA through the DH domain. A) HDX-MS Woods’ plots showing individual peptides across PRG (left) or RhoA (right) plotted by the difference in deuteration of these regions within the GS-DH/PH–RhoA complex as compared to the DH/PH–RhoA complex after 10,000s of deuterium exposure. Blue lines represent regions with less deuteration or solvent accessibility in the presence of the GS, while red lines represent regions with more deuteration in the presence of the GS. The areas of the plot highlighted in light blue are hotspots for GS-induced decreases in deuteration. B) Crystal structure of GS-DH/PH–RhoA (PDB: 5JHG) colored by HDX-MS difference data after 10,000s of exposure. The GS is colored pink. The eyeball icons indicate the viewing angles that produced the images of the interface between GS, DH, and RhoA switch I (1) and the interface between RhoA switch II and the DH domain (2). The distance between the centers of mass of PRG GS (712-728) and RhoA Switch II (57-78) is shown in (2).

### Initial cryo-EM reconstruction of the PRG DH/PH– RhoA complex

To delve further into the details of this allosteric mechanism, we used single-particle cryo-EM to pursue a high-resolution structure of PRG DH/PH–RhoA to compare to the available crystal structures of GS-DH/PH–RhoA. We collected micrographs of DH/PH– RhoA using a 200 keV Glacios to ascertain if cryo-EM could be used to resolve this ∼65 kD, small particle. High-contrast particles resembling the protein complex were clearly visible on the micrographs (Figure S6A and B), and after pre-processing, particle picking, and extraction, 2D classification showed class averages with high-resolution features and representing diverse viewing angles (Figure S6C). Despite this, 3D ab-initio reconstruction and subsequent refinement appeared to suffer from poor orientation distribution and anisotropy (Figures S5A, S6D and S8). At the same time, Kim et al. published findings detailing an alternative 3D reconstruction strategy, named high-resolution homogenous ab-initio reconstruction (HR-HAIR)^24^, which obtains initial particle orientations during ab-initio reconstruction exclusively using high spatial frequency information and iterating over numerous rounds through very small Fourier radius steps to improve 3D reconstruction for small particles. This strategy also entails 2D classification parameters for optimal sorting of small particles before ab-initio. We re-processed our dataset using these parameters in cryoSPARC (see Experimental Procedures) and obtained 2D class averages with clearer high-resolution detail and representing a variety of viewing angles. The resulting 3D reconstruction had more defined secondary structure features accompanied by a drastic improvement in the cFAR score (Figures S7B, E, and S8). These trends support that particle images simply underwent poor alignment, leading to our previous substandard reconstructions and low cFAR scores. As part of the HR-HAIR pipeline, local refinement is run after ab-initio reconstruction in order to preserve the output 3D orientation assignments, which are not carried forward in other cryoSPARC refinement methods. However, subsequent local refinement of our higher quality reconstruction from HR-HAIR yielded limited improvements and sometimes led to a noisier, poorer quality map than the input reconstruction. As we suspected that this was still due to inaccuracies in alignments likely owing to the small size of the protein, we next sought to improve the signal-to-noise in our data.

### Higher resolution cryo-EM reconstruction of ∼46 kD of the PRG DH/PH–RhoA complex

In pursuit of a more interpretable reconstruction, we collected an additional dataset using a 300 keV Krios equipped with an energy filter to help reduce background and improve the signal-to-noise ratio of our small particl es. With the previously utilized HR-HAIR processing parameters, we obtained 2D class averages with even higher resolution features (Figure S11C) and a 3D reconstruction with more structural detail than those from our Glacios dataset. We also optimized HR-HAIR parameters for our sample by further decreasing the ab initio Fourier radius step to 0.001 from 0.005, resulting in inconsistent mild improvements (Figure S9). Since we had observed that local refinement of reconstructions from our Glacios data led to minimally improved, if not poorer quality, maps, we next wanted to optimize local refinement to further improve particle orientations after HR-HAIR. We noticed that local refinement using a Gaussian blurred mask, which is common practice^26^, produced a reconstruction rife with discontinuities in the protein backbone (Figure S10A–B). Instead, we dilated and minimally padded the PRG DH–RhoA HR-HAIR reconstruction in cryoSPARC Volume Tools and used this as a tight mask in local refinement. The resulting final 3D reconstruction, representing ∼46 kD of the PRG DH/PH–RhoA complex, at sub-4 Å resolution displayed improved interpretability, backbone continuity, and side chain density (Figures 4A and S11D). Importantly, the corrected FSC curve closely tracked the tight-mask curve, indicating the lack of appreciable tight masking artifacts (Figure S11F).

**Figure 4.**
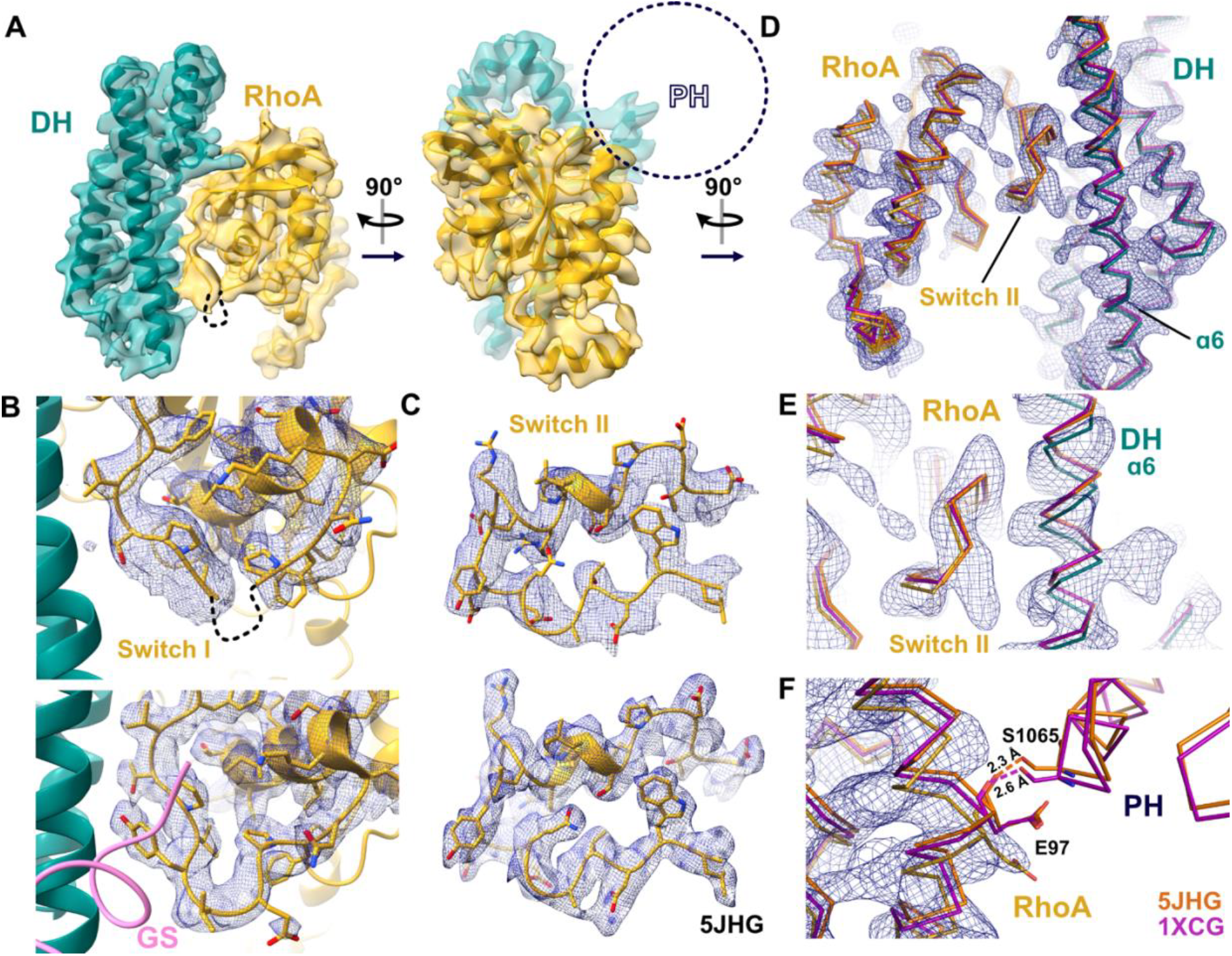
DH/PH–RhoA cryo-EM reconstruction and comparison with the GS-DH/PH–RhoA crystal structure. A) Cryo-EM structure of 46 kD of the PRG DH/PH–RhoA complex. Density for PRG DH (teal) and RhoA (yellow) are clearly visible, but density for the PH domain is completely missing. B) Close-up views of RhoA Switch I in the DH/PH–RhoA cryo-EM structure (top) and in a previous crystal structure of GS-DH/PH–RhoA (5JHG; bottom) shown with electron density contoured at 1 σ. Switch I is poorly resolved in our cryo-EM structure lacking the GS but well-resolved in the crystal structure containing the GS. C) Close-up views of RhoA Switch II in the DH/PH–RhoA cryo-EM structure (top) and in the GS-DH/PH–RhoA crystal structure (5JHG; bottom) shown with electron density contoured at 1 σ. Switch II is well-resolved in both DH/PH–RhoA and GS-DH/PH– RhoA structures. D) Ribbon representation of GS-DH/PH–RhoA crystal structures 5JHG (orange) and 1XCG (purple) superimposed on our DH/PH–RhoA cryo-EM structure and map. Residues 873-890 of the DH α5 helix were aligned in this and the two following panels. Close-up views of E) RhoA Switch II and PRG DH α6 and F) RhoA Glu97 and PRG PH Ser1065, which form a hydrogen bond in the crystal structures. Shifts in α-helices within PRG and RhoA are observed in our cryo-EM structure lacking the GS relative to the crystal structures that contain the GS.

### Cryo-EM data support an allosteric mechanism for the activity-enhancing effect of the GS

Our final reconstruction, at an estimated global resolution of ∼3.7 Å, shows the PRG DH domain bound to RhoA but lacks any density for the PH domain. Although the PH domain was clearly visible in a scarce number of 2D class averages (Figure S6E), there was no trace of it in any of our 3D reconstructions no matter how we attempted to sort the particles (Figure 4A). This was unexpected, as the PH domain was unambiguously accounted for in our low-resolution SEC-SAXS data, but also because the PH domain is well-resolved in crystal structures of GS-DH/PH– RhoA^16,27,28^ (Figures 4A and 1B). Within RhoA, the tip of the Switch I loop (Glu32 through Tyr34) is unstructured, and the backbone density C-terminal to the tip is poorly defined and likely dynamic relative to the rest of RhoA (Figure 4B), differing from the crystal structure of GS-DH/PH–RhoA in which Switch I is well-resolved (Figure 4B). This corroborates our HDX-MS data demonstrating that the GS stabilizes Switch I. In contrast, Switch II density is well-defined, similar to in the presence of the GS (Figure 4C). This was surprising in light of our HDX-MS data showing that Switch II displayed a more drastic deuteration decrease than Switch I in the presence of the GS (Figure 3A). To further investigate this apparent discrepancy and the missing PH domain, we obtained a low-resolution cryo-EM reconstruction of GS-DH/PH–RhoA to compare with our DH/PH–RhoA cryo-EM reconstruction (Figure S13). Again, density for the DH domain and RhoA is well-defined (Figure S13C). In addition, extra density is present in a location that would correspond to part of the PH domain. The appearance of this extra density in the presence of the GS is striking, as we were unable to recover any density for the PH domain in our DH/PH–RhoA reconstructions regardless of any particle sorting or refinement strategies tested. Interestingly, this extra density shields Switch II from the solvent, reconciling the deuteration decrease observed in Switch II in the presence of the GS in our HDX-MS data with its defined density in both our DH/PH–RhoA cryo-EM structure and the crystal structures containing the GS. In other words, the increased Switch II protection in the presence of the GS may be due to decreased dynamics of the PH and its corresponding association with RhoA. In support of this, the PRG DH α6 helix is shifted along its axis toward its N-terminus in our DH/PH–RhoA cryo-EM structure relative to two crystal structures of GS-DH/PH–RhoA^16,27^, which would position the PH differently relative to the rest of the complex (Figure 4D and E). In our DH/PH–RhoA structure, we also noticed a shift in the RhoA helix containing residues 90-102 away from the PH domain (Figure 4F). This alpha helix includes Glu97, which forms a hydrogen bond with Ser 1065 in PRG PH in the GS-DH/PH–RhoA crystal structures^16,27^, indicating that RhoA may shift depending on association with the PH as a result of allosteric effects from the GS. Overall, our data illustrate that the GS allosterically stabilizes the PRG catalytic core–RhoA complex, primarily RhoA Switch I and PRG PH, to enhance PRG activity towards RhoA (Figure 5).

**Figure 5.**
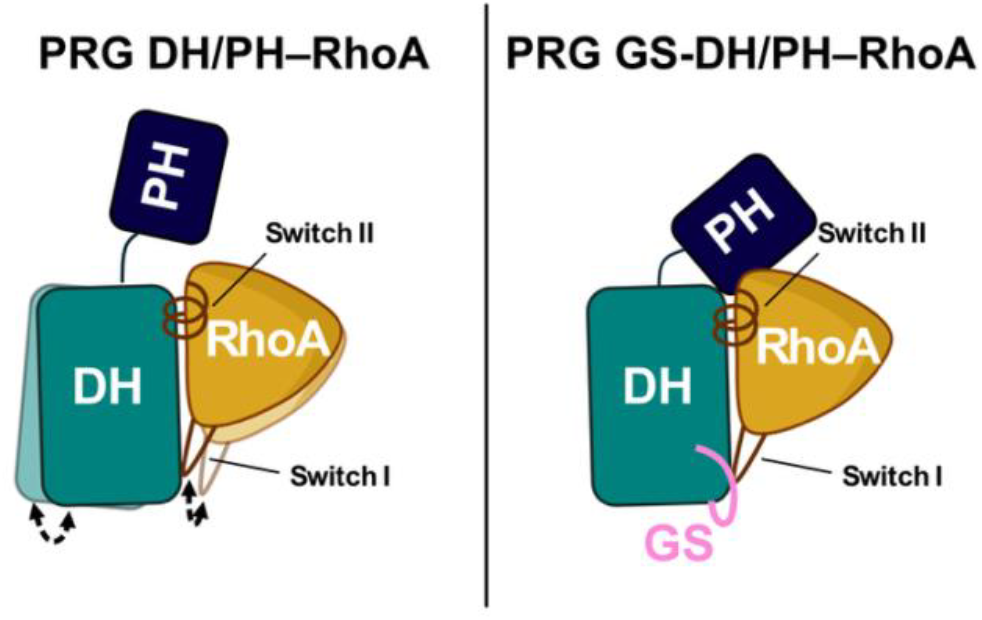
Proposed model for the GEF Switch-mediated enhancement of PRG activity. In the PRG DH/PH–RhoA complex lacking the GS, DH, PH, and RhoA are relatively dynamic. Switch I of RhoA, which constitutes part of the guanine-nucleotide binding pocket, shows particularly high dynamics in the absence of the GS. In the GS-DH/PH–RhoA complex, the GS interacts directly with DH, remodeling it such that the PH domain becomes more stably associated with the complex, and the switch regions of RhoA become less dynamic, promoting GEF activity toward RhoA.

## Discussion

Here, we present integrative structural information demonstrating that the GS potentiates PRG activity likely by stabilizing the complex between PRG and nucleotide-free RhoA, the catalytic intermediate for guanine nucleotide exchange^29^. Our biochemical assays demonstrate that interactions with the DH α1 helix and α2-α3 loop are required for the GS to augment PRG activity. Our HDX-MS and cryo-EM data reveal that within this intermediate complex, the GS stabilizes the DH domain and likely the PH domain in addition to Switch I of RhoA. These data inform the mechanism by which the GS enhances PRG activity, encompassing the GS-proximal interactions required for this function and the allosteric effects on the rest of the PRG–RhoA complex. To our knowledge, this is the first presentation of a mechanism for the GS in any RhoGEF.

The GS packs against the PRG DH α1 helix and α2-α3 loop to stabilize a conformation that potentiates RhoA activation. Previous observations in LARG and p115-RhoGEF revealed some of the structural requirements for GS function, including the importance of the tryptophan in the center of the GS hydrophobic pocket (LARG Trp769; p115-RhoGEF Trp398; PRG Trp716) and of GS interactions with the α1 helix through a Glu790 in LARG DH (Glu737 in PRG DH)^22,23^. Our study expands upon these structural requirements by showing that in addition to GS–α1 interactions, GS–α2-α3 loop interactions are equally important for GS function. These findings also illustrate the inherently dynamic nature of the DH domain and suggest a model in which its structural stabilization can amplify RhoGEF activity. This mirrors how the PH domain of PRG enhances PRG activity potentially through stabilization of the DH α6 helix^16^. Conceivably, this structural attribute of the DH domain could be exploited by other activators.

RhoGEFs catalyze guanine nucleotide exchange on Rho GTPases by facilitating the release of GDP, the rate-limiting step in the reaction. Here, the GS stabilizes the PRG catalytic core–nucleotide-free RhoA complex, a critical reaction intermediate. The switch regions of RhoA undergo conformational changes in transition from GDP-bound to nucleotide-free during guanine nucleotide exchange^17,30^. In the GDP-bound state, RhoA Switch I directly engages the guanine ring and cooperates with Switch II to coordinate the Mg^2+^ ion that bridges the β-phosphate of GDP to RhoA^31^. RhoGEFs pry Switches I and II away from the nucleotide binding pocket, disrupting the pocket and the Mg^2+^ coordination network, encouraging divalent cation release along with GDP^22^. Switch I of RhoA is partially disordered in our DH/PH–RhoA cryo-EM structure, and the GS-mediated stabilization of Switches I and II supported by our HDX-MS data connects the GS enhancement function to the guanine nucleotide exchange mechanism. While the GS likely stabilizes Switch I in part by packing against it, the Switch II region is solvent-protected through a long-range allosteric mechanism. Overall, the GS appears to enhance PRG activity, at least in part, by conferring stabilization of these RhoA regions to promote the release of GDP from RhoA.

The GS stabilizes Switch I through direct packing, as observed in crystal structures of the GS-DH/PH– RhoA complex^16,27^. Additionally, the GS interacts with Ile733 and Glu737 of the DH α1 helix, which likely stabilizes the helix in an optimal conformation for contact with RhoA Switch I. This is supported by our cryo-EM data of the PRG DH/PH–RhoA complex; in the absence of the GS, the complex lacks strong density for a short segment of Switch I (Figure 4B). This segment includes Tyr34, which packs against the DH α1 helix in GS-DH/PH–RhoA crystal structures^16,27^. By contrast, the nature of the allosteric mechanism for Switch II stabilization remains unclear. In the DH/PH– RhoA cryo-EM structure, Switch II is well-resolved even in the absence of the GS. A low-resolution cryo-EM reconstruction of GS-DH/PH–RhoA exhibits partial density for the PH domain, unlike DH/PH–RhoA, indicating that the GS also stabilizes the PH domain. Our DH/PH–RhoA cryo-EM structure gives insight into how the GS does this as seen in the shift in the DH α6 helix relative to GS-containing crystal structures (Figure 4E). Taken together with the α2-α3 loop stabilization observed in our HDX-MS data, we propose a two-pronged allosteric GS mechanism. First, the GS packs against the DH α1 helix and stabilizes it into a conformation that is optimal for packing against RhoA Switch I. Through a second route, the GS stabilizes the DH α2-α3 loop, causing shifts in the α6 helix that propagate to stabilize the PH domain in a complex-bound state in which it caps a region of RhoA that includes Switch II.

The position of the PH domain in our GS-DH/PH– RhoA cryo-EM reconstruction generally corroborates the crystal structures^16,27^ (Figure S13). However, it is worth noting that the PH domain forms only a small interface with RhoA in these crystal structures (Figure 4E) and is otherwise stabilized by crystal contacts with a neighboring PH domain through a sticky hydrophobic surface along β5, β6, and β7 strands, a crystal contact commonly observed in RhoGEF crystal structures^22,23,27,32–34^. This raises the question of how closely these crystal structures encapture the native state of the PH domain and its interactions, highlighting that other structural techniques should be used to confirm interactions involving the PH domain, such as cryo-EM.

In this study, we were able to determine a sub-4 Å structure of a smaller than 50 kD particle using cryo-EM without the use of fiducial markers. Our ability to determine this cryo-EM structure depended upon formation of a stable protein complex and rigorous sample quality validation through SEC, SAXS, and DSF showing that the complex was monodisperse, stable, non-aggregated, and at least mostly globular (Figures 2A-E and S2A-D). Above all, robust biochemistry remains fundamental for the structural determination of small particles using cryo-EM. Advances in cryo-EM technology, such as energy filters, and data processing through HR-HAIR have also helped to enable structure determination of these previously intractable, small targets. We found that HR-HAIR^24^ can be complemented by a tighter masking strategy during subsequent local refinement and that HR-HAIR parameters should be optimized on a sample-to-sample basis. Even still, 3D alignment for these small particles remains error-prone, as indicated by the moderate resolution of our reconstructions, highlighting the need for further advances in data processing or cryo-EM technology.

In summary, the GS plays a significant role in potentiating RhoA activation by PRG and other GS-containing RhoGEFs. Our results provide a foundation for understanding the structural basis of the fully active state of PRG and potentially other GS-containing, RhoA-activating RhoGEFs. The significance of the GS as a shared feature among a wide number of other RhoGEFs is unknown, warranting further investigation of the GS in other RhoGEFs (Figure S1). In crystal structures of p115-RhoGEF, density for the GS is lost upon addition of an N-terminal autoinhibitory sequence accompanied by a decrease in p115-RhoGEF activity, revealing GS destabilization as an underlying mechanism for autoinhibition. However, how this autoinhibition is relieved by activators remains unexplained. In addition, whether the GS of PRG is subject to a similar “on-off” switching mechanism also remains unclear. Future studies focusing on the role of the GS in PRG regulation in the context of the full-length protein in conjunction with other regulatory accessory regions and binding partners will be pivotal in understanding how PRG is regulated and inform strategies for exploiting these mechanisms for the development of inhibitors to treat PRG-driven pathologies.

### Experimental procedures

#### GEF Switch Sequence Alignment

RhoGEF amino acid sequences containing a tryptophan residue within 25 residues N-terminal to the UniProt-annotated DH domain were identified as candidates for GS-containing RhoGEFs. These sequences were aligned using ClustalOmega^35^ and visualized using ESPript^36^.

#### Plasmids and Cloning

A plasmid expressing human RhoA, pLV_DORA_RhoA_IRES, was generated by the Hayer lab and gifted to us through Dr. Sean Collins^37^. DNA encoding RhoA without the CAAX lipidation site (residues 1-189) was cloned into a modified pMal expression vector (pMALc2H_10_T) through Gibson Assembly, resulting in a soluble RhoA construct with a TEV-cleavable N-terminal MBP-10xHis tag^22^. Human pDONR221 PRG cDNA was obtained courtesy of the Center for Personalized Diagnostics through DNASU (HsCD00860223)^38^. DNA encoding PRG DH/PH (residues 729-1081) was cloned into the above described pMALc2H_10_T vector, and PRG GS-DH/PH (residues 712-1081) was generated by insertion of a nucleotide sequence corresponding to residues 712-728 into this construct through a previously described site-directed mutagenesis method^39^. This method was also used to generate pMALc2H_10_T PRG PH (residues 927-1081) through deletion of the DH domain coding sequence from the PRG DH/PH construct. PRG GS-DH/PH point mutants were generated using the same site-directed mutagenesis method.

#### Protein Expression and Purification

Soluble RhoA was co-purified with GDP using a similar procedure as previously described for Rac1 and Cdc42, with some differences^34^. Namely, the lysis buffer included a different protease inhibitor cocktail: 1 μM bestatin, 0.124 μM aprotinin, 1 μM trans-Epoxysuccinyl-L-leucylamido(4-guanidino)butane (E-64), 6.8 μM leupeptin, and 0.15 μM soybean trypsin inhibitor. Size-exclusion chromatography was performed using a Superdex 200 Increase 10/300 GL (Cytiva).

PRG DH/PH and GS-DH/PH were purified as previously described for DH/PH^40^. To form PRG–RhoA complexes, RhoA was added at a molar ratio of 1:1 to the flow-through from the Ni-NTA resin containing PRG GS-DH/PH or DH/PH and incubated for 15 minutes at 4°C. EDTA was then added to 10 mM to strip Mg^2+^ and nucleotide from RhoA to stabilize the complex, and this was incubated overnight at 4°C. PRG or PRG–RhoA complexes were supplemented with 2% glycerol, concentrated using a 10kDa cutoff concentrator, and loaded on an ENrich™ SEC 70 10 x 300 column (Bio-Rad) equilibrated with 20 mM HEPES pH 8, 200 mM NaCl, 2 mM DTT, 2% glycerol, and 2 mM EDTA. If necessary, fractions containing PRG or PRG–RhoA complexes were concentrated using a 10 kDa MWCO Amicon concentrator, and purified protein was immediately used for experiments.

The isolated PRG PH domain was expressed in Rosetta (DE3) pLysS *E. coli* cells, which were harvested, lysed, and clarified in the same manner as DH/PH^40^. Clarified PRG PH lysate was incubated with Ni-NTA resin for 1 hour. Resin was washed with Buffer A (20 mM HEPES pH 8, 300 mM NaCl, 2 mM DTT) followed by Buffer A with 10 mM imidazole then eluted with Buffer A with 250 mM imidazole. Elutions were dialyzed into 20 mM HEPES pH 7, 100 mM NaCl, 2 mM DTT and treated with TEV protease at a 1:16 molar ratio TEV:MBP-His-PH overnight to remove the N-terminal MBP-His tag. The dialyzed sample was supplemented to 20 mM imidazole and 300 mM NaCl and incubated with Ni-NTA resin for 1 hour to capture the cleaved tag. Flow-through was processed on an ENrich SEC70 (BioRad) SEC column in 20 mM HEPES, pH 7, 200 mM NaCl, 2 mM DTT. Protein was flash frozen, stored at -80 °C, and thawed when protein was needed for assays.

#### GTPase Exchange Activation Assay

The guanine nucleotide exchange activity of PRG was measured by a FRET-based mant-GTP exchange assay^41^. Unprenylated RhoA·GDP was mixed at a concentration of 2 μM with 0.8μM N-methyl-anthraniloyl-GTP (mant-GTP; Abcam), a fluorescent, non-hydrolyzable GTP analog, in buffer containing 20 mM HEPES pH 8, 100 mM NaCl, 5 mM MgCl_2_, 1 mM DTT, and 5% glycerol and incubated at room temperature for 30 minutes until 10 nM PRG was added to initiate the reactions. Reactions were dispensed in triplicate onto a 384-well black plate and inserted into a SpectraMax® M5 (Molecular Devices) microplate reader. Fluorescence at Ex = 280 nm, Em = 450 nm was measured over the course of 10 minutes in 30 second intervals at room temperature (24 °C). The reaction traces were fitted to a straight line in Prism, and the slope of the line was calculated. Each independent experiment was performed in technical triplicate with PRG GS-DH/PH WT and at least one mutant. The slope for each mutant technical replicate was normalized to the mean of the GS-DH/PH WT slopes for each independent experiment. For each independent experiment, the normalized technical triplicate slopes for each variant were averaged to produce a single normalized biological replicate activity value that was plotted and used for further statistical analysis.

#### Differential Scanning Fluorimetry

Protein was brought up to 96.7 μl using DSF Buffer (20 mM HEPES pH 8, 200mM NaCl, 2mM DTT) before 3.3 μL 300x SYPRO Orange or Prolite Orange was added. Reactions were aliquotted in 30 μl technical triplicates into 0.2 ml PCR tubes. Reactions were spun down before being placed into a BioRad CFX96™ System configured using BioRad Maestro to increase sample temperature from 22 °C up to 95 °C at a rate of 1 °C/min. Output was read in FRET mode with each well of interest designated as an Unknown with a FRET fluorophore in Channel 6, according to manufacturer’s specifications. Raw melting curves were normalized as percentages in Prism with the lowest value in each curve set to 0 and the highest value set to 100. Normalized curves were truncated after the 100% value and fit to a Boltzmann sigmoidal curve. Inflection points were calculated from each curve and reported as the melting temperature. Independent experiments were performed in technical triplicate. Technical replicate values were averaged to obtain melting temperature values for each biological replicate.

#### Size-Exclusion Chromatography Small-Angle X-ray Scattering

Small-angle X-ray scattering (SAXS) experiments were performed at the SIBYLS beamline 12.3.1 at the Advanced Light Source. This beamline has additional inline instrumentation and detectors coupled to a size-exclusion column^42,43^. DH/PH–RhoA and GS-DH/PH– RhoA were purified and concentrated in SEC running buffer (20mM HEPES pH7, 200mM NaCl, 2mM DTT, 2% glycerol, 2mM EDTA). The X-ray wavelength was set to 1.127 Å and the sample-to-detector distance to 2,100 mm. This combination gives scattering vectors (q) ranging from 0.01 Å^-1^ to 0.4 Å^-1^. The scattering vector is q = 4πsinθ/λ, where 2θ is the scattering angle. The SAXS flow cell was coupled to an Agilent 1260 Infinity HPLC system using a Shodex PROTEIN KW-802.5 8 x 300mm SEC column equilibrated with running buffer at a flow rate of 0.65 ml min^-1^. The eluent was subject to the following: 1) ultra-violet light at 280 nm, 2) multi-angle light scattering, 3) quasi-elastic light scattering, 4) SAXS, and 5) a refractometer. For SAXS measurements, 2 sec X-ray exposures were collected continuously during a 25 min elution and recorded using a Pilatus3 2M pixel array detector. All frames for analyses had one SAXS frame corresponding to the running buffer before the detection of a peak subtracted from each.

The radius of gyration (Rg) was calculated for each of the subtracted frames using the Guinier approximation: I(q) = I(0) exp(™q^2^Rg^2^/3) with the limits qRg < 1.3. The elution peak was compared to the integral of ratios to background and Rg relative to the recorded frame using the program RAW^44^. Uniform Rg values across an elution peak represent a homogeneous sample. Final merged SAXS profiles, derived by integrating multiple frames at the elution peak, were used for further analyses. The Guinier plot was calculated to provide information on the aggregation state, the volume of correlation (Vc) to estimate the molecular weight^45^, and the pair distribution function [P(r)] to calculate the maximal inter-particle dimension^46^.

*Hydrogen-Deuterium Exchange Mass Spectrometry* Prior to conducting hydrogen/deuterium exchange experiments, optimal quench conditions were determined to generate the best enzymatic peptide coverage map for PRG DH/PH, GS-DH/PH, and RhoA, as previously described^47,48^. Briefly, 3 μl purified PRG DH/PH–RhoA (2.7 mg/ml in 20 mM HEPES pH 8, 200 mM NaCl, 2 mM DTT, 2mM EDTA, 2% glycerol) or GS-DH/PH–RhoA (4.5 mg/ml in 20 mM HEPES pH 8, 200 mM NaCl, 2 mM DTT, 2mM EDTA, 2% glycerol) were diluted with 9 μl of H_2_O buffer at 0 °C, and then mixed with 18 μl of ice cold quench buffers containing 0.1 M glycine (pH 2.4), 16.6% glycerol and varying concentration of GuHCl (0.08, 0.8, 1.6 and 3.2 M). The quenched samples were subjected to on-line proteolysis and LC/MS analysis. The best sequence coverages of PRG DH/PH, GS-DH/PH, and RhoA were obtained using the 0.08 M GuHCl quench buffer.

PRG–RhoA exchange stock solutions (23 μM of each PRG DH/PH–RhoA and PRG GS-DH/PH–RhoA) were prepared in 8.3 mM Tris, 150 mM NaCl pH 7.2 and kept on ice. HDX experiments were initiated by adding 50 μl of exchange stock solutions to 150 μl of D_2_O buffer (8.3 mM Tris, 150 mM NaCl, pDread7.2) and incubating for various times intervals (10, 100, 1000, 10,000 and 100,000 sec) at 0 °C. At each time, 36 μl of exchange mixture was withdrawn and quenched with 54 μl of ice-cold quench buffer (0.1 M glycine pH 2.4, 0.08 M GuHCl). Quenched samples were aliquoted and immediately frozen on dry ice. Non-deuterated and equilibrium deuterated control samples were also prepared for back exchange correction^49^.

All frozen samples were later loaded onto a cryogenic autosampler and automatically thawed at 4 °C before digestion on an immobilized pepsin column (16 μl bed volume) at 25 μl/min. Proteolytic fragments were collected on a trap column and separated on an Acclaim PepMap RSLC C18 reverse phase analytical column (ThermoSci, 0.3 × 50 mm, 2 μm, 100 Å) using a linear acetonitrile gradient (5%™45% over 30 min). The effluent was directed into an OrbiTrap Elite Mass Spectrometer (ThermoFisher Scientific, San Jose, CA). Instrument settings were optimized to minimize the back-exchange^50^. The data were acquired in either MS1 profile mode or data-dependent MS/MS mode. Peptide identification was done with the aid of Proteome Discoverer software (ThermoFisher). The centroids of the mass envelopes of deuterated peptides were calculated with HDExaminer (Sierra Analytics Inc, Modesto, CA) and then converted to corresponding deuterium incorporation with corrections for back-exchange^51^.

#### Cryo-EM Grid Preparation and Data Acquisition

For the PRG DH/PH–RhoA Glacios dataset, complex was prepared at 0.3 mg/ml and mixed with 0.08 mM DDM before applying 3 μL of the solution to a glow discharged 300-mesh Quantifoil (1.2/1.3) grid in a Vitrobot (Thermo Fisher Scientific) set to 4 °C and 100% humidity. The loaded grid was blotted for 4 seconds at a blot force of 20 before being plunge-frozen into ethane cooled by liquid nitrogen. Data were collected at the Biological Electron Microscopy (BioEM) Facility at the University of California, Davis on a Glacios cryogenic transmission electron microscope (Thermo Fisher Scientific) equipped with a Gatan K3 direct electron detector using SerialEM. HR-HAIR and non-HR-HAIR workflows utilized the same subset of 5,406 movies, but the non-HR-HAIR workflow included an additional 2,022 movies (Figure S5).

For the PRG DH/PH–RhoA Krios dataset, grids were prepared in the same manner as above, except PRG DH/PH–RhoA was mixed with 0.02 mM lubrol and applied to a glow discharged 300-mesh UltrAuFoil (1.2/1.3) grid. Data were collected at the Pacific Northwest Center for Cryo-EM (PNCC) on a Krios cryogenic transmission electron microscope (Thermo Fisher Scientific) equipped with a Gatan K3 direct electron detector and a BioContinuum HD energy filter using SerialEM. For the PRG GS-DH/PH–RhoA Glacios dataset, grids were prepared in the same manner as the PRG DH/PH–RhoA Krios Dataset. Data were collected at the BioEM Facility at the University of California, Davis.

#### Cryo-EM Data Processing

All cryo-EM data processing steps were performed using cryoSPARC. All movies underwent pre-processing steps of motion correction, patch CTF estimation, and micrograph denoising before particle picking and extraction.

The Glacios DH/PH–RhoA particles were processed through two rounds of 2D classification, ab-initio reconstruction, and non-uniform refinement, resulting in a 3D reconstruction completely lacking the PH domain (Figure S5A). Subsequent local refinement was performed with a DH–RhoA mask and Gaussian priors in an attempt to prevent severe misalignment of the low signal-to-noise particles during 3D refinement. Details for data processing are shown in Figure S5. A subset of these particles were alternatively processed using high resolution homogeneous ab-initio reconstruction (HR-HAIR)^24^ to improve particle alignment (Figure S5B). A single round of 2D classification with HR-HAIR parameters was sufficient to produce detailed 2D class averages (Figure S7). Particles selected from 2D classification were sorted into three classes using HR-HAIR ab-initio, which produced a junk class, a lower quality, opposite-handed class, and a higher quality, correct-handed class. The correct-handed class was chosen for subsequent local refinement with a DH–RhoA mask, strict Gaussian priors, and search extents to prevent severe misalignment in 3D refinement. The local refinement map showed improvement compared to the non-HR-HAIR map but only marginal improvement compared to the initial ab-initio HR-HAIR map (Figure S8).

The Krios DH/PH–RhoA particles were cropped to 1 Å/px during extraction followed by 2D classification using HR-HAIR parameters. Details for data processing are shown in Figure S9. Resulting 2D class averages were biased towards a single view of the protein complex, motivating the rebalancing of the 2D classes in cryoSPARC with a rebalance factor of 0.8. With a more view-balanced particle stack, initial particle orientations were determined using HR-HAIR with a smaller Fourier radius step than previously published^24^, which produced minor improvements in the map. Particles were uncropped and then processed in 3D using another round of HR-HAIR and local refinement with a tight DH–RhoA mask that was generated from the HR-HAIR volume, dilated and minimally padded using Volume Tools in cryoSPARC. The resulting map was of high enough quality to build a model into but was an excessively large file that hindered convenient file handling. As such, the particles that were used for the local refinement were cropped to half of the super-resolution pixel size, re-extracted, and re-processed using local refinement using a DH–RhoA mask that was generated using a similar approach as the previous mask. This process resulted in our final DH– RhoA map, which was sharpened using Phenix before being used for model building.

#### Model Building, Refinement, and Validation

Using the AlphaFold Server^52^, a model of the PRG DH– RhoA complex was generated using the sequences of the DH domain and RhoA constructs used in our study. The highest scoring model was fit into our cryo-EM map using ChimeraX^53^. Parts of the model that were completely outside of map density were deleted in Coot 0.9.8.95^54^, and then the model was run through rigid body real-space refinement in Phenix 1.21.2-5419^55^ after adding hydrogens and with minimization_global enabled^56^ using the unsharpened map. After this, the model was further modified and rebuilt into the sharpened map in Coot using tight geometric restraints. The map is mostly isotropic with some less interpretable regions where the density is smeared between secondary structure features. Density modification using EMReady improved interpretability in some of these areas, and this output map was used to help model building where needed^57^. Crystal structures of PRG GS-DH/PH–RhoA (1XCG, 3T06, 5JHG, and 5JHH) were also used to assist decision making during model building. Some structural elements in both the DH domain and RhoA clearly needed to be shifted to better match the density (Fig. 4D-F). After building was complete, regions of the structure that formed helices or β-sheets were manually confirmed and annotated, and Phenix real-space refinement was run using secondary structure restraints. Model validation was performed in Phenix^56^(Table 2 & Fig. S12B). EMRinger was also run within Phenix^58^. The associated web interface was used to run the 3D FSC program to calculate sphericity and generate a 3D FSC histogram and directional FSC plot^59,60^. The Q score was calculated during the validation stage of data deposition^61^. Structure figures were generated using ChimeraX and PyMOL^62^. Some software was accessed through membership in the SBGrid Consortium^63^.

**Table 2.**
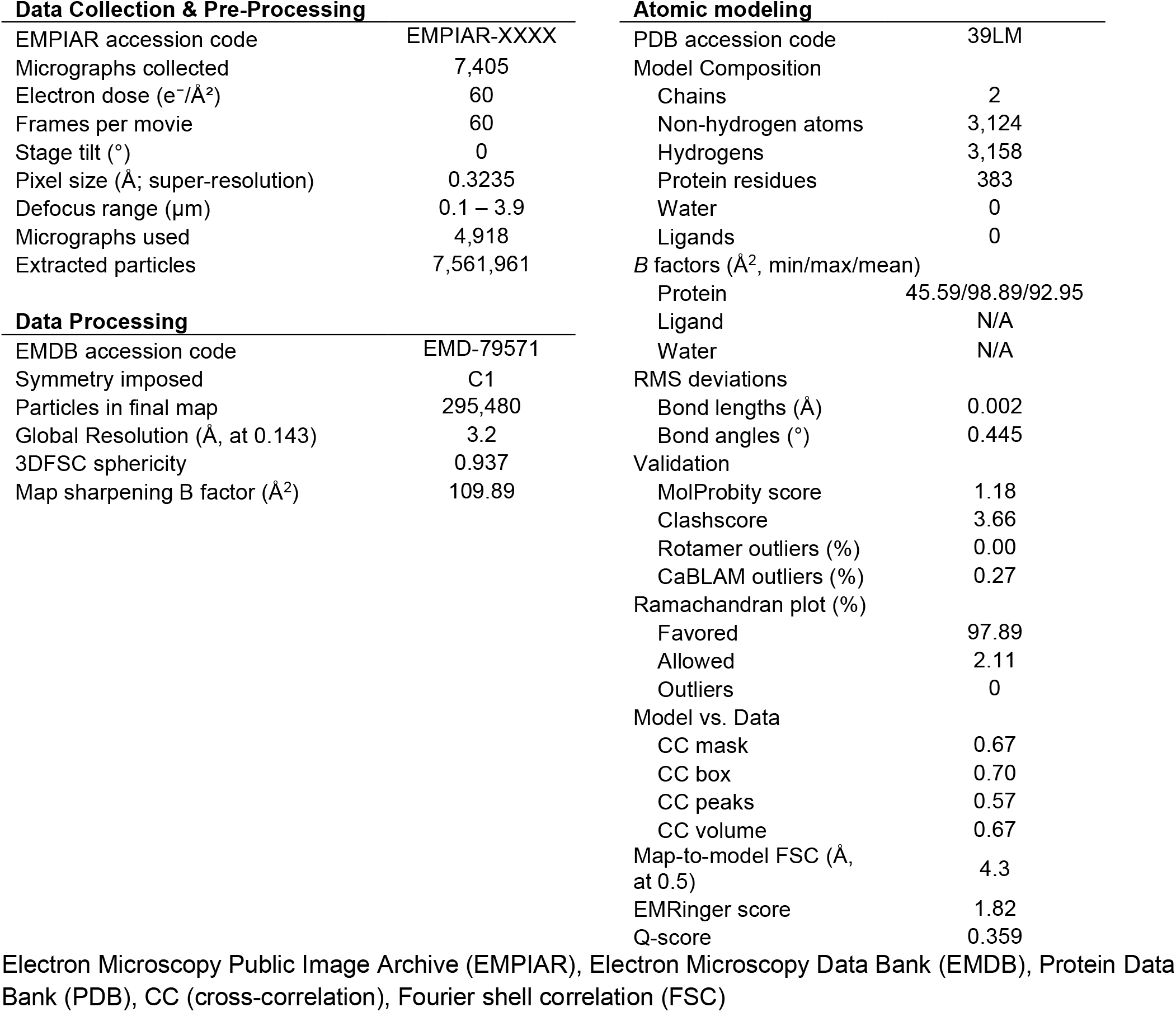
Cryo-EM Data Collection, Processing, and Modeling Statistics.

#### Statistical Analysis

All statistical analyses were performed using GraphPad Prism from at least three independent biological replicates, each performed in technical triplicate. For the mant-GTP exchange assay, PRG GS-DH/PH mutants were evaluated for statistical significance using a one-sample t-test comparing the mutant normalized activity biological replicate values to a theoretical WT GS-DH/PH activity value of 1 followed by a Holm-Šídák correction for multiple comparisons (*p < 0.05). For the GS-DH/PH mutant differential scanning fluorimetry assay, melting temperature biological replicate values were compared to GS-DH/PH WT melting temperature biological replicate values. Statistical significance was determined using one-way ANOVA with a post hoc Dunnett’s test for multiple comparisons, using WT as the control (*p < 0.05, **** p<0.0001).

## Supporting information

This article contains supporting information.

Document S1. Table S1 and Figures S1–13

Data S1. PRG DH/PH–RhoA and PRG GS-DH/PH–

RhoA HDX-MS profiles

## Resource Availability

### Lead contact

Requests for further information and resources should be directed to and will be fulfilled by the lead contact, Jennifer Cash.

### Data availability

The PRG DH/PH–RhoA EM data can be accessed as EMDB entry EMD-79571 and EMPIAR entry EMPIAR-XXXX. The model has been deposited as PDB entry 39LM. The SAXS data for PRG GS-DH/PH–RhoA and DH/PH–RhoA can be accessed as SASDB entry XXXX and XXXX, respectively.

### Materials availability

All unique/stable reagents generated in this study are available from the lead contact with a completed materials transfer agreement.

## Acknowledgements

We would like to thank Michal Hammel at the Lawrence Berkeley National Laboratory for assistance with SEC-SAXS data interpretation. We would also like to thank Fei Guo for support in the UC Davis BioEM facility and Hemang Patel and Camille Scott for IT and HPC support for cryo-EM data storage and processing. We would like to thank Marcelo de Farias at the Pacific Northwest Cryo-EM Center for assisting with Krios cryo-EM data collection. We furthermore would like to thank Emma Barpal for providing technical support for the project. Finally, we would like to thank members of the Cash Lab for feedback on the manuscript.

## Author contributions

E.B. cloned PRG GS-DH/PH variants, expressed these variants, performed some of their purifications and ran the corresponding biochemical assays along with M.V.B. S.L. performed HDX-MS experiments on purified PRG–RhoA samples and analysis of the raw data. M.V.B. performed all other benchwork, including preparing samples for HDX-MS and SEC-SAXS experiments and preparing and screening samples for cryo-EM, processing cryo-EM data, analyzing experimental results, preparing figures, and writing the initial draft of the manuscript. J.N.C. provided guidance on cryo-EM data processing and built the PRG DH/PH– RhoA model.

## CRediT author contributions

Conceptualization - M.B. and J.N.C.

Data curation - M.B., E.B., J.N.C., S.L.

Formal analysis - M.B., E.B., J.N.C., S.L.

Funding acquisition - M.B. and J.N.C.

Investigation - all authors

Methodology - M.B. and J.N.C.

Project administration - J.N.C.

Resources - J.N.C.

Supervision - M.B. and J.N.C.

Validation - M.B. and J.N.C.

Visualization - M.B. and J.N.C.

Writing - original draft - M.B and J.N.C.

Writing - review & editing - all authors

## Funding and additional information

Research reported in this publication was supported by the National Institute of General Medical Sciences of the National Institutes of Health under Award Number R35GM146664 (to J.N.C.). This work was supported in part by the UC Davis Comprehensive Cancer Center (P30CA093373; to J.N.C.). This work was supported by NIH Training Grant T32 GM 007377 (to M.V.B). A portion of this research was supported by NIH grant R24GM154185 and performed at the Pacific Northwest Center for Cryo-EM (PNCC) with assistance from Marcelo De Farias. SAXS data was collected at the Advanced Light Source (ALS) at the SIBYLS beamline, a national user facility operated by Lawrence Berkeley National Laboratory on behalf of the Department of Energy, Office of Basic Energy Sciences, through the Integrated Diffraction Analysis Technologies (IDAT) program, supported by DOE Office of Biological and Environmental Research. Additional support comes from the National Institute of Health project ALS-ENABLE (P30 GM124169). Molecular graphics and analyses performed with UCSF ChimeraX, developed by the Resource for Biocomputing, Visualization, and Informatics at the University of California, San Francisco, with support from National Institutes of Health R01-GM129325 and the Office of Cyber Infrastructure and Computational Biology, National Institute of Allergy and Infectious Diseases. The content is solely the responsibility of the authors and does not necessarily represent the official views of the National Institutes of Health.

## Abbreviations and nomenclature

(RhoGEF): Rho guanine-nucleotide exchange factor
(GDP): guanosine diphosphate
(GTP): guanosine triphosphate
(Dbl): diffuse B-cell lymphoma
(DH): Dbl homology
(PH): pleckstrin homology
(PDZ): Postsynaptic density protein 95, Discs large protein, and Zonula occludens-1
(cryo-EM): cryo-electron microscopy
(HDX-MS): hydrogen-deuterium exchange mass spectrometry
(SEC-SAXS): small-angle X-ray scattering
(RMSD): root mean squared deviation
(SDS-PAGE): sodium dodecyl sulfate–polyacrylamide gel electrophoresis
(mant-GTP): N-methyl-anthraniloyl-GTP
(P(r)): pair distance distribution
(EOM): employed ensemble optimization method
(WT): wild-type
(MBP): maltose binding protein
(GST): glutathione S-transferase
(DTT): dithiothreitol
(HEPES): 4-(2-hydroxyethyl)-1-piperazineethanesulfonic acid
(NaCl): sodium chloride
(EDTA): ethylenediaminetetraacetic acid
(CV): column volumes
(IPTG): isopropyl β-D-1-thiogalactopyranoside
(MWCO): molecular weight cutoff
(DDM): n-Dodecyl-Beta-Maltoside
(PNCC): Pacific Northwest Center for Cryo-EM
(CTF): contrast transfer function
(FSC): fourier shell correlation
(GuHCl): guanidine hydrochloride
(LC/MS): liquid chromatography/mass spectrometry
(D_2_O): deuterium oxide
(Tris): Tris(hydroxymethyl)aminomethane
(MgCl_2_): Magnesium chloride
(Rg): radius of gyration
(Vc): volume of correlation

