## Supplemental Information for "Cryo-EM structure of the sub-50 kD PDZ-RhoGEF catalytic core–RhoA complex reveals an allosteric mechanism for GEF switch"

**Supplemental Information Table S1 and Figures S1-S13**

**Table S1. DSF melting temperatures for PRG GS-DH/PH variants**

| Variant | $T_m$ (°C) ± 1 SD |
|---|---|
| WT | 44.3 ± 1.0 |
| W716F | 38.8 ± 1.0 |
| E737A | 41.9 ± 0.8 |
| I733S | 32.3 ± 1.8 |
| DH/PH | 34.9 ± 0.2 |

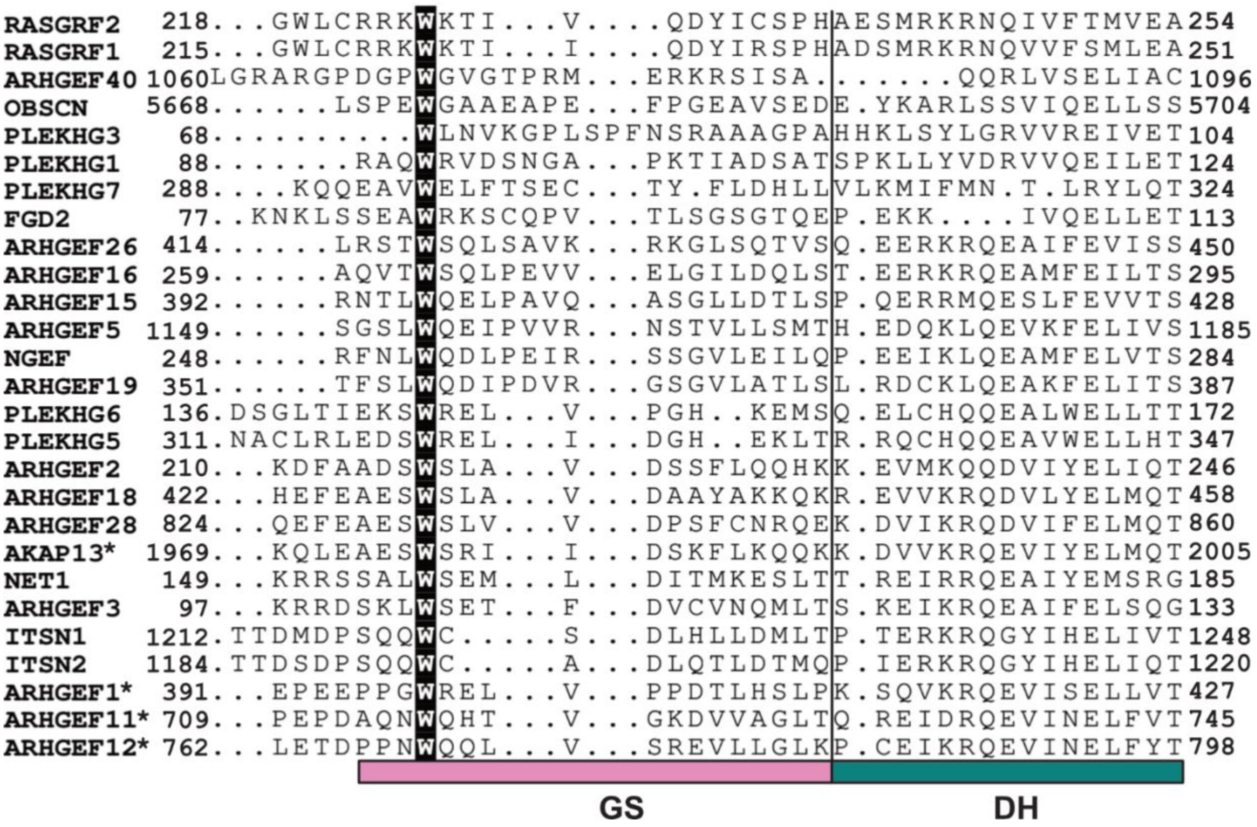


**Figure S1. Potential GEF switch-containing RhoGEFs.** RhoGEFs containing at least one tryptophan (highlighted in black) within 25 residues N-terminal to the UniProt-annotated DH domain were considered as candidates for GS-containing RhoGEFs and were aligned using ClustalOmega and visualized using ESPript. Asterisks next to the names indicate RhoGEFs in which the GS fold was observed in at least one published structure.

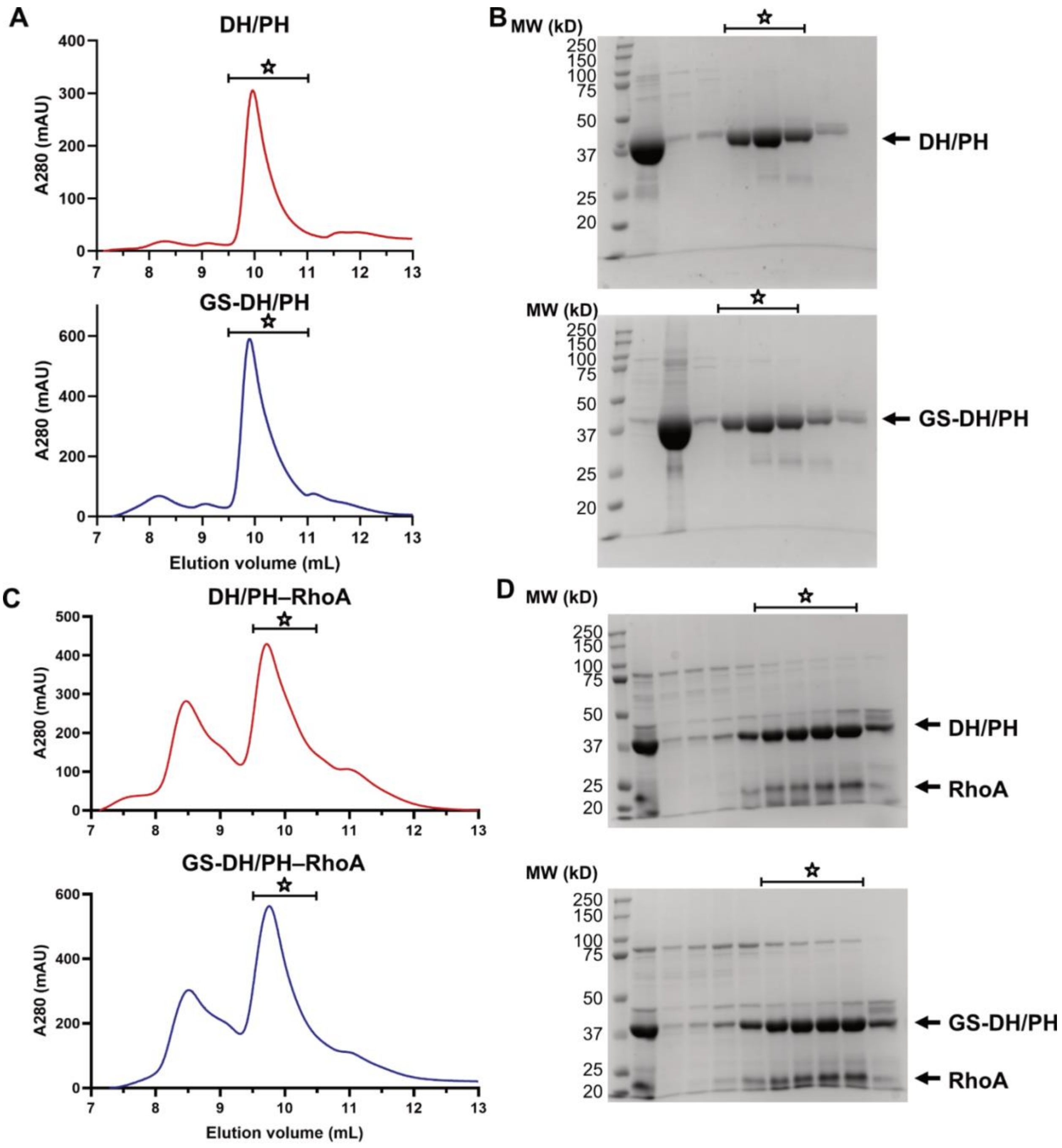


**Figure S2. PRG DH/PH and GS-DH/PH purification.** A) Size-exclusion chromatography traces and B) SDS-PAGE gels for DH/PH (top) and GS-DH/PH (bottom). C) SEC traces and D) SDS-PAGE gels for DH/PH–RhoA (top) and GS-DH/PH–RhoA (bottom). Stars indicate regions of the SEC peak or gel containing protein or protein complex.

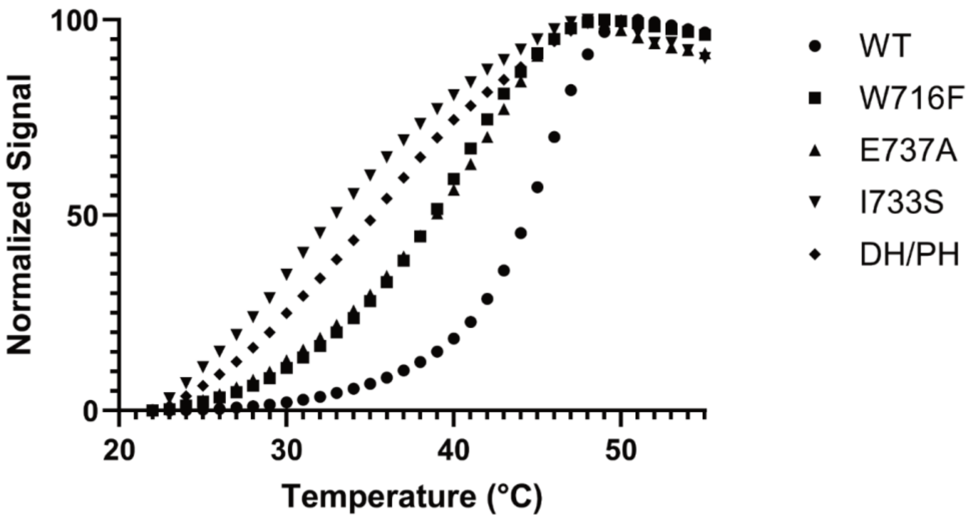


**Figure S3. Sample quality of PRG GS-DH/PH variants.** Representative DSF normalized melting curves from PRG GS-DH/PH variants and DH/PH (see Figure 1D).

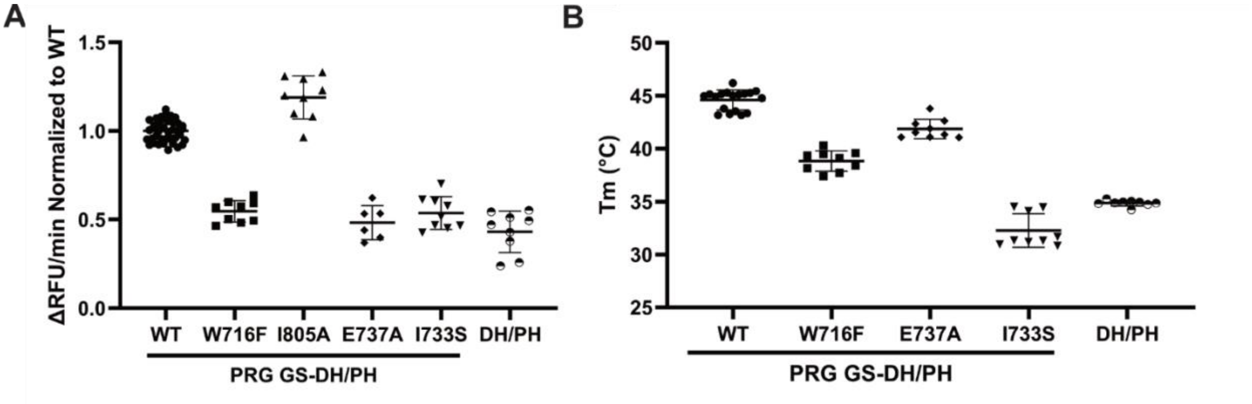


**Figure S4. Mant-GTP exchange and DSF assays with pooled technical replicates.** A) mant-GTP exchange assay showing the normalized activities of PRG DH/PH and GS-DH/PH variants towards RhoA. Individual points represent technical replicates from multiple independent experiments. Raw ΔRFU/min values were normalized to the mean of the WT ΔRFU/min values. Error bars represent ±1 SD. B) DSF assay comparing the melting temperatures of PRG DH/PH and GS-DH/PH variants. Individual points represent technical replicates from multiple independent experiments. Error bars represent ±1 SD.

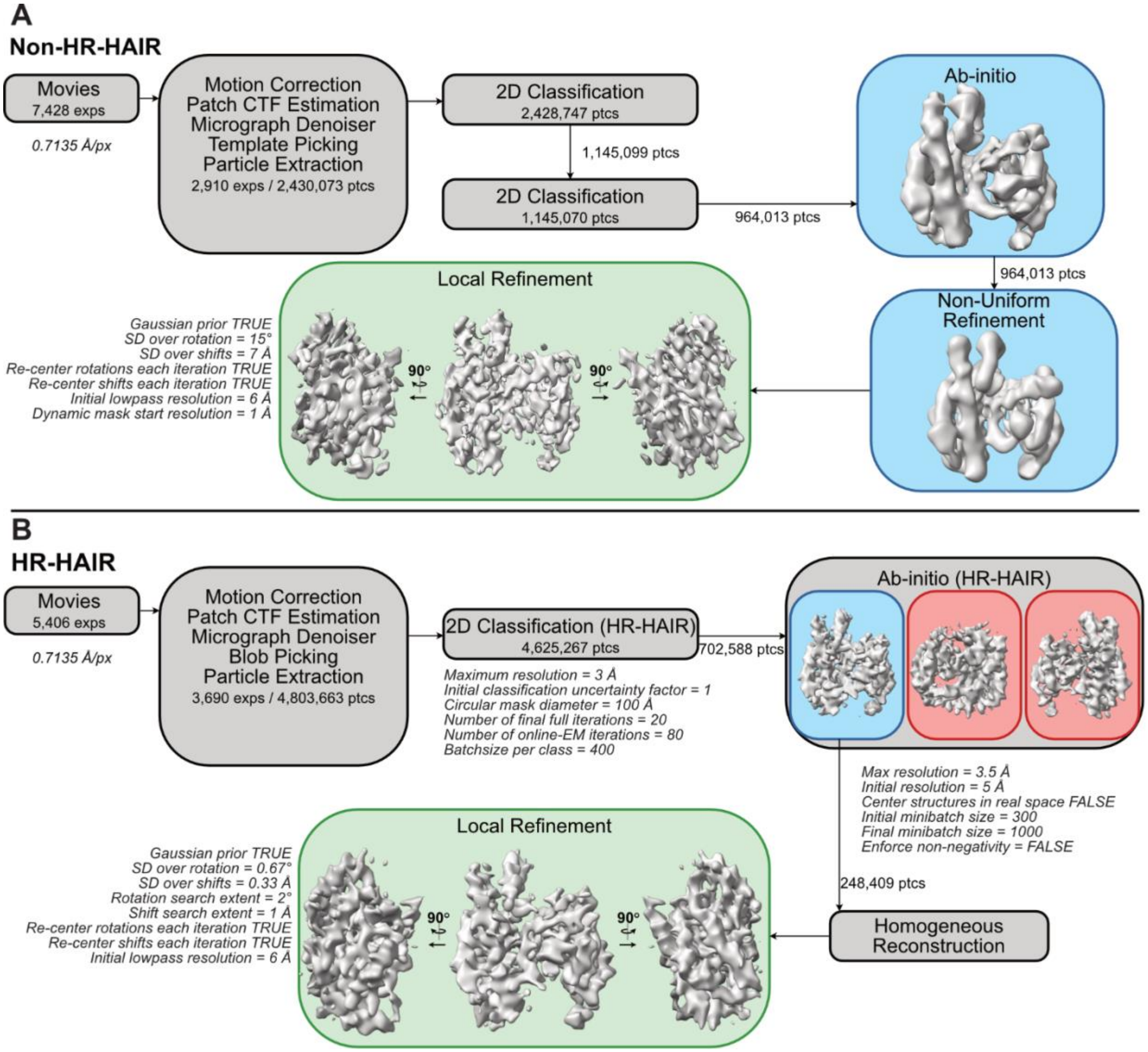


**Figure S5. DH/PH–RhoA Glacios Workflow.** A) Non-HR-HAIR data processing pathway for DH/PH–RhoA. B) HR-HAIR data processing pathway for DH/PH–RhoA. Intermediate 3D reconstructions are highlighted in blue, and final 3D reconstruction is highlighted in green. Unused 3D reconstructions are highlighted in red. All processing steps were performed in cryoSPARC. Processing parameters that deviated from default values for 2D classification, 3D reconstruction, and 3D refinement jobs are included in the workflow diagram.

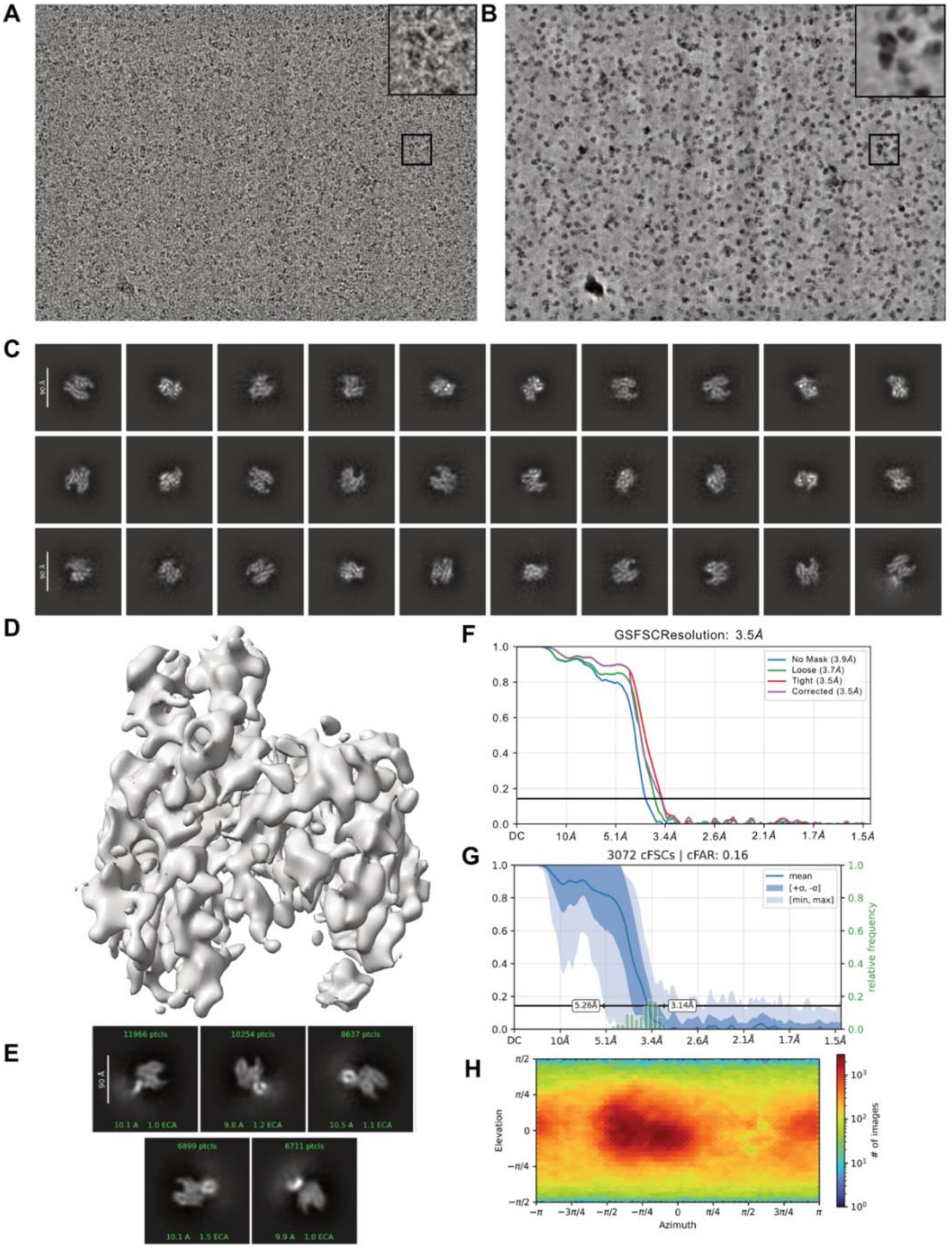


**Figure S6. PRG DH/PH–RhoA cryo-EM reconstruction from data collected on Glacios.** A) Data collection and processing of PRG DH/PH–RhoA showing a representative raw micrograph of PRG DH/PH–RhoA and B) denoised image of this micrograph, each with a zoomed in view shown in inset, C) 2D class averages, and D) 3D reconstruction. E) A small subset of 2D class averages showing the PH domain as a bright white spot that is circular or smeared. Despite this mass clearly representing the PH domain in 2D, there was no indication of its presence in any of our 3D reconstructions. Diagnostic plots generated in cryoSPARC showing the F) gold standard FSC curve, G) conical FSC plot with cFAR score, and H) viewing direction distribution plot.

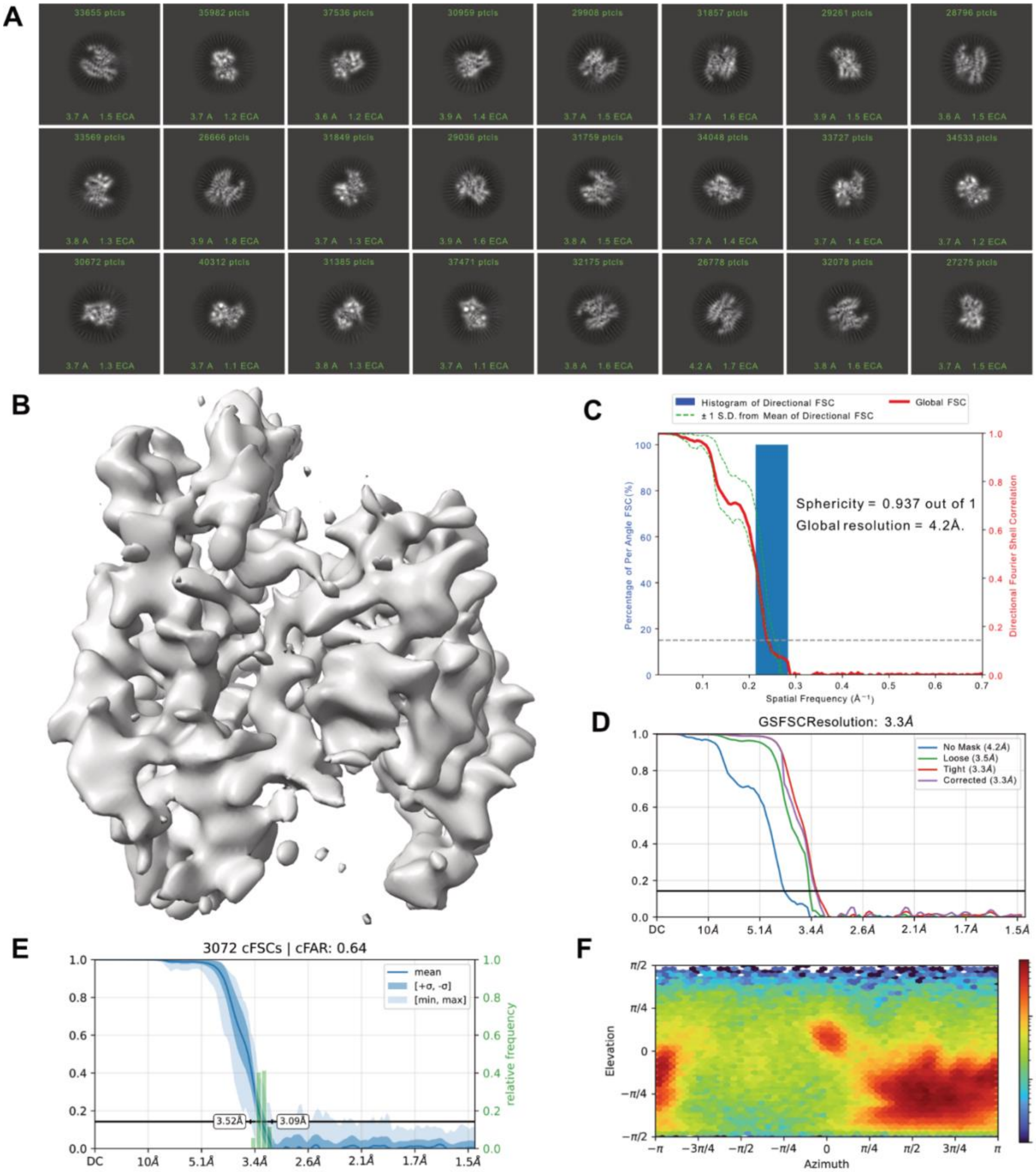


**Figure S7. PRG DH/PH–RhoA cryo-EM reconstruction from data collected on Glacios and processed using HR-HAIR.** PRG DH/PH–RhoA A) 2D class averages and B) 3D reconstruction after local refinement, generated using HR-HAIR parameters. C) 3DFSC histogram and directional FSC curve calculated by the 3DFSC program (https://3dfsc.salk.edu/). Diagnostic plots generated in cryoSPARC showing the D) gold standard FSC curve, E) conical FSC plot with cFAR score, and F) viewing direction distribution plot.

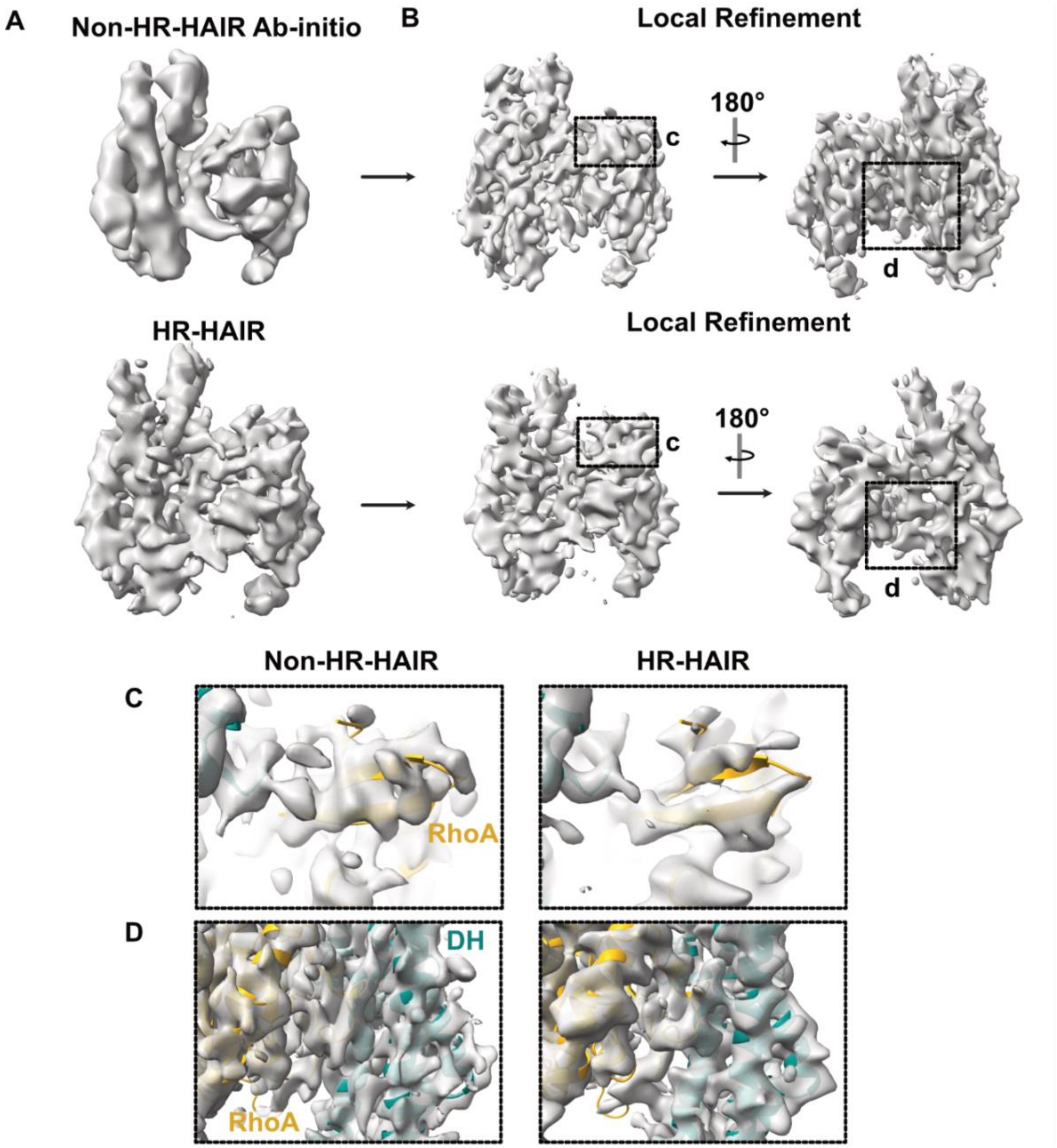


**Figure S8. Comparison of results from HR-HAIR and non-HR-HAIR processing pipelines for DH/PH–RhoA data.** A) Comparison between ab-initio reconstruction without HR-HAIR (top, see Figure S5A) and with HR-HAIR parameters (bottom, see Figure S5B). B) Comparison between local refinement after standard ab-initio reconstruction (top) and after HR-HAIR (bottom), featuring close-up views of C) two β-strands in RhoA and D) a portion of the RhoA–DH interface in the non-HR-HAIR pipeline (left) and the HR-HAIR pipeline (right). The HR-HAIR data processing workflow resulted in clearer secondary structure density and reduced streakiness.

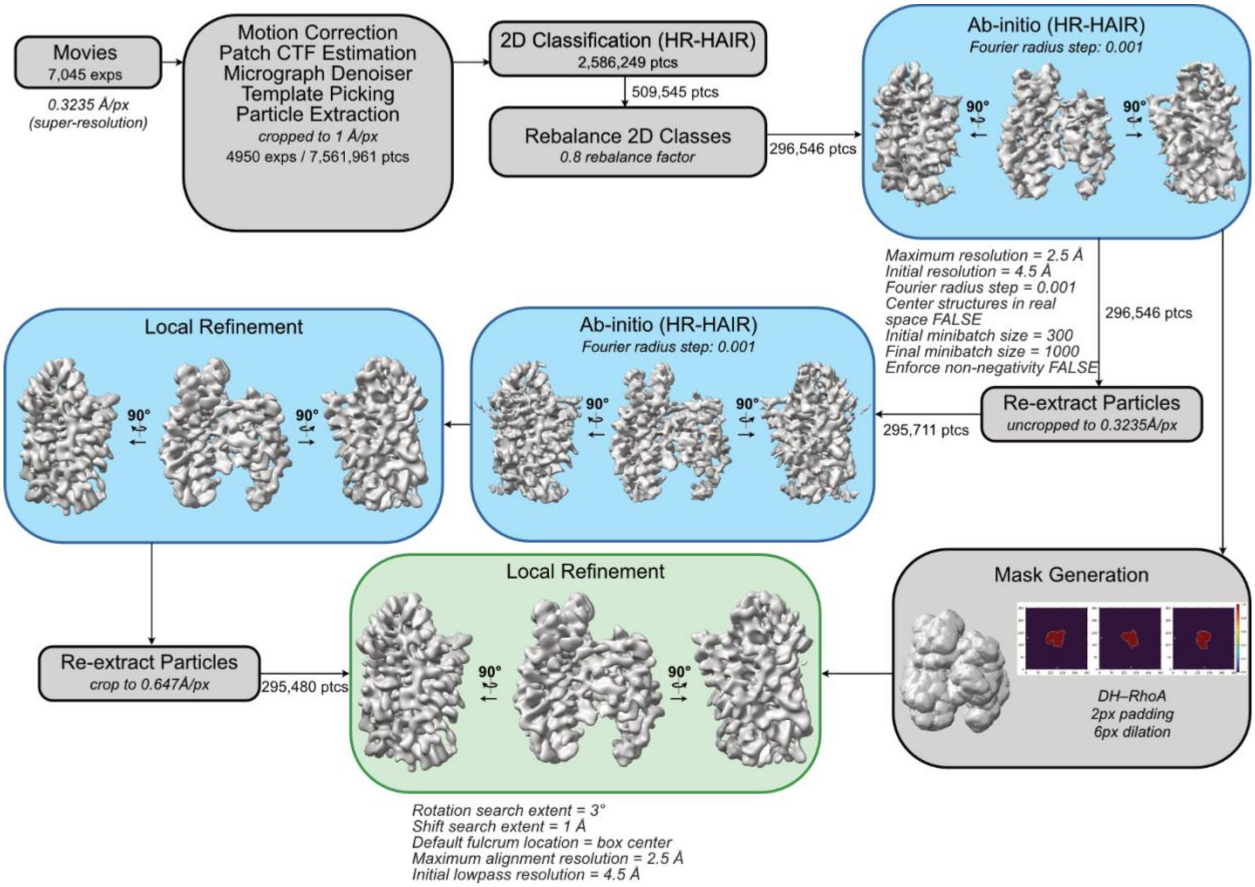


**Figure S9. Data processing workflow for DH/PH–RhoA cryo-EM data collected on Krios.** Intermediate 3D reconstructions are highlighted in blue, and final 3D reconstruction is highlighted in green. All processing steps were performed in cryoSPARC. Processing parameters that deviated from default values for 2D classification, 3D reconstruction, and 3D refinement jobs are included in the workflow diagram.

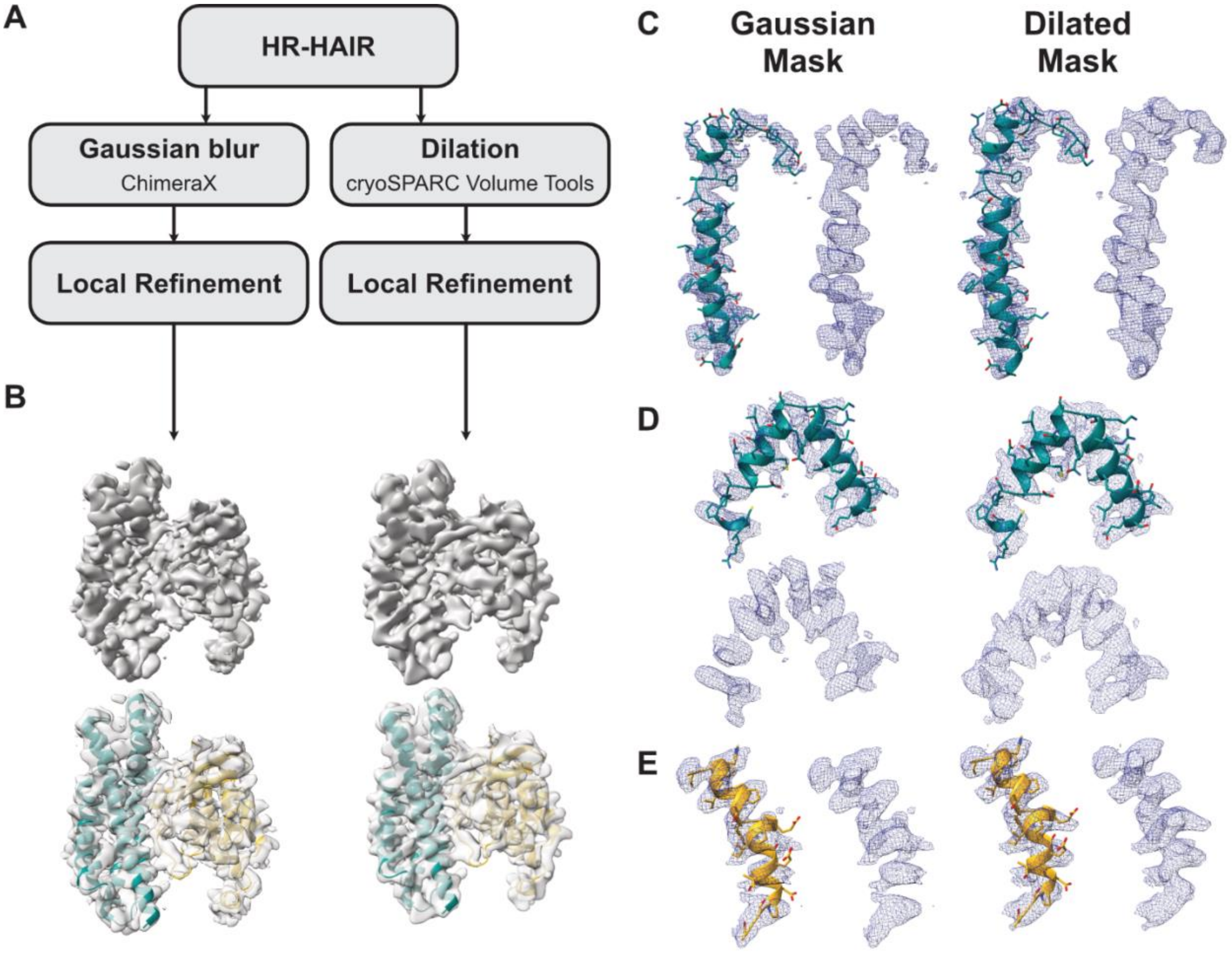


**Figure S10. Effect of tighter masking through dilation on local refinement.** A) Mask generation strategies used for DH/PH–RhoA data processing, resulting in B) 3D reconstructions. Models of C) DH residues 766–803, D) DH residues 835–867, E) and RhoA residues 87–107 fit into densities generated by local refinement using a Gaussian-blurred mask as compared to a non-Gaussian-blurred, dilated mask.

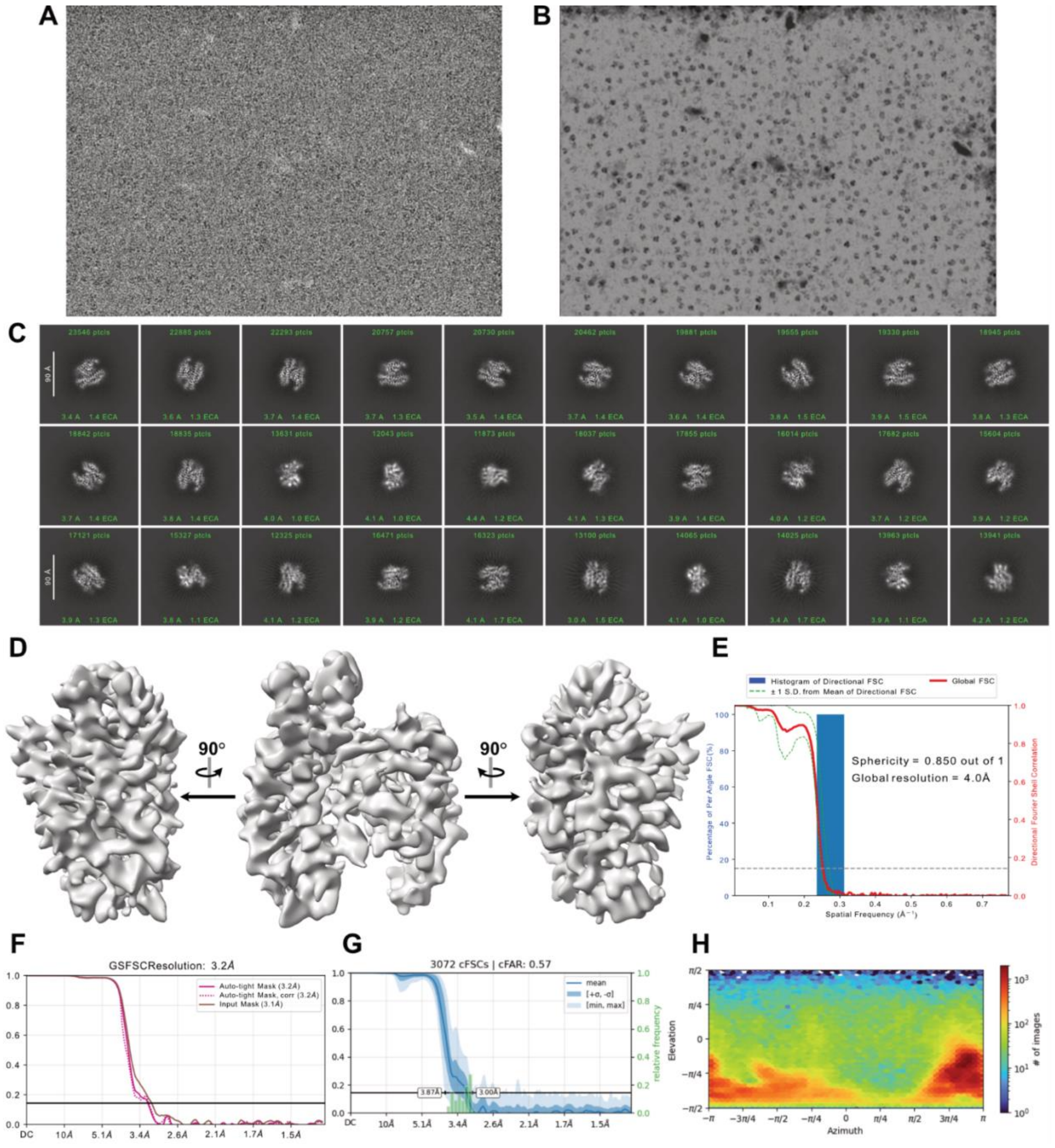


**Figure S11. PRG DH/PH–RhoA cryo-EM reconstruction from data collected with Krios.** A) Representative raw micrograph, B) denoised image of this micrograph, C) 2D class averages, and D) 3D reconstruction of PRG DH/PH–RhoA. E) 3DFSC histogram and directional FSC curve calculated by the 3DFSC program (https://3dfsc.salk.edu/). Diagnostic plots generated in cryoSPARC showing the F) gold standard FSC curve, G) conical FSC plot with cFAR score, and H) viewing direction distribution plot.

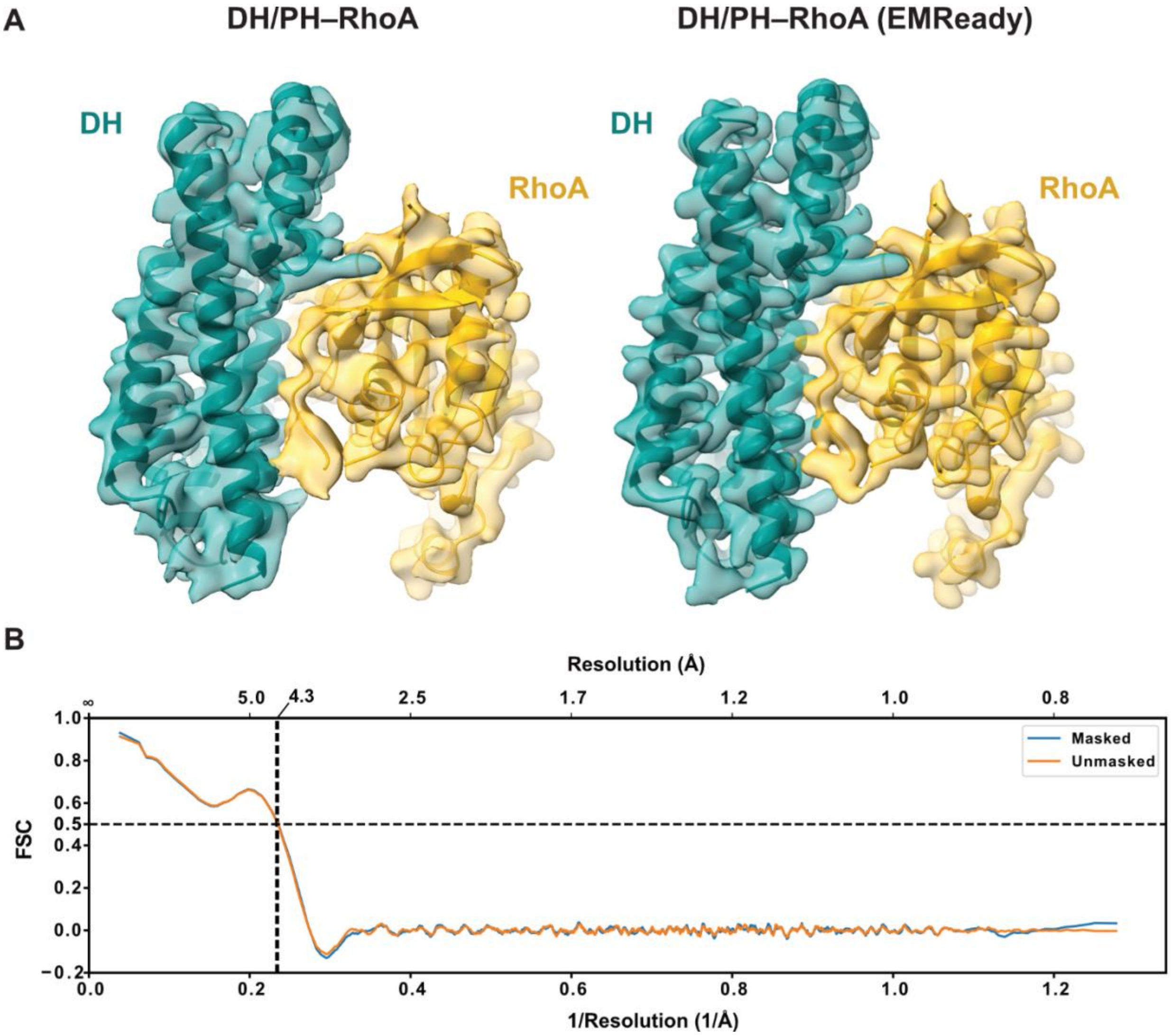


**Figure S12. Cryo-EM map enhancement through EMReady to guide modeling.** A) DH/PH–RhoA sharpened cryo-EM map (left) and DH/PH–RhoA map enhanced using EMReady (right). B) Model-to-map fit of our DH/PH–RhoA cryo-EM structure.

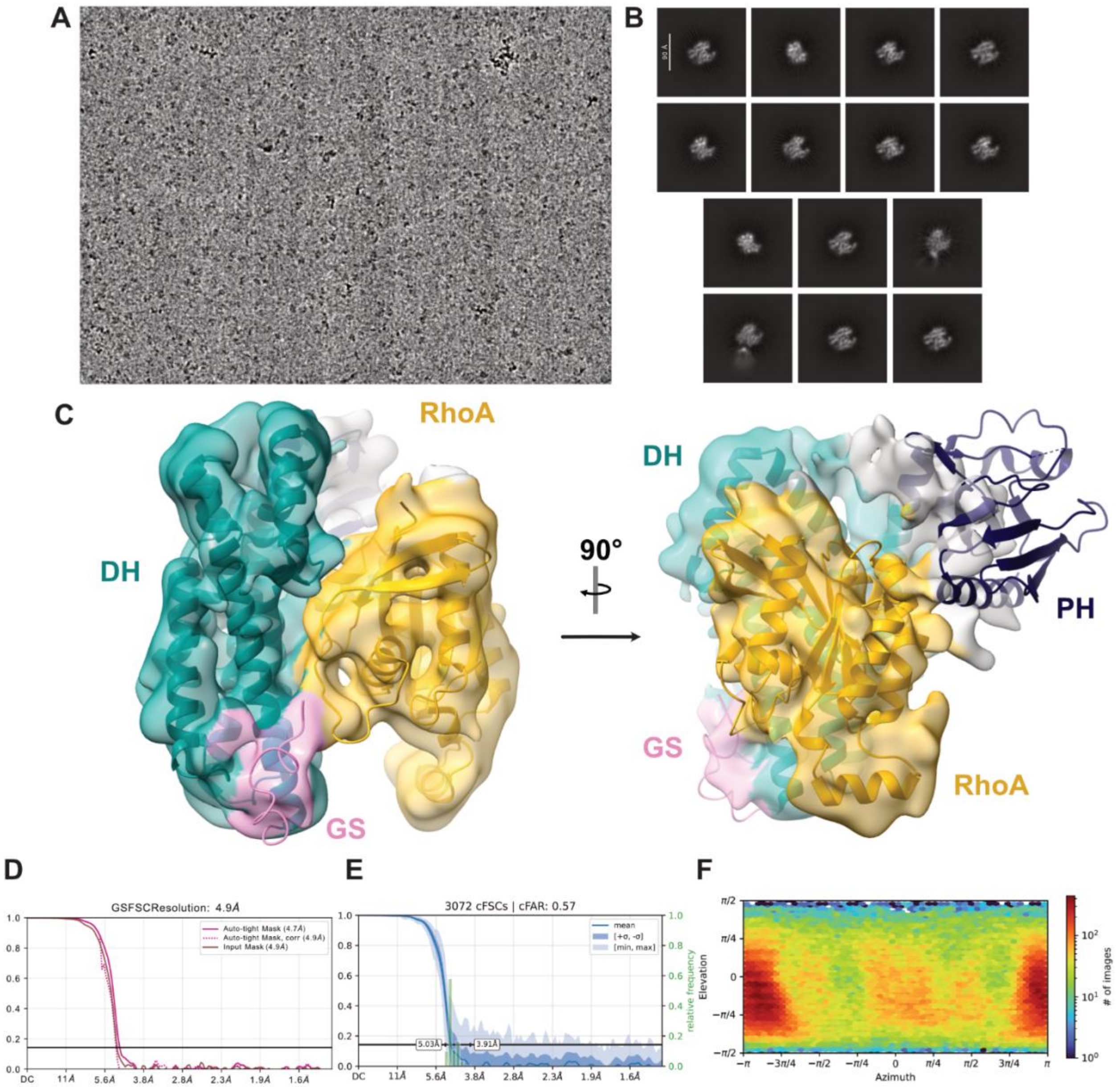


**Figure S13. Low resolution cryo-EM reconstruction of PRG GS-DH/PH–RhoA.** A) Representative micrograph and B) 2D class averages of PRG GS-DH/PH. C) Low resolution 3D reconstruction of PRG GS-DH/PH–RhoA with rigid body fitted crystal structure 5JHG. This reconstruction contains density that likely corresponds to the PH domain, based on its location. Diagnostic plots generated in cryoSPARC showing the D) gold standard FSC curve, E) conical FSC plot with cFAR score, and F) viewing direction distribution plot.
